# The canonical E2Fs together with RETINOBLASTOMA-RELATED are required to establish quiescence during plant development

**DOI:** 10.1101/2022.12.05.519120

**Authors:** Magdolna Gombos, Cécile Raynaud, Yuji Nomoto, Eszter Molnár, Rim Brik-Chaouche, Hirotomo Takatsuka, Ahmad Zaki, Dóri Bernula, David Latrasse, Keito Mineta, Fruzsina Nagy, Xiaoning He, Hidekazu Iwakawa, Erika Őszi, Jing An, Takamasa Suzuki, Csaba Papdi, Clara Bergis, Moussa Benhamed, László Bögre, Masaki Ito, Zoltán Magyar

## Abstract

Maintaining stable and transient quiescence in differentiated and stem cells, respectively, requires repression of the cell cycle. The plant RETINOBLASTOMA-RELATED (RBR) has been implicated in stem cell maintenance, presumably by forming repressor complexes with E2F transcription factors. Surprisingly we find that mutations in all three canonical E2Fs do not compromise the cell cycle, but similarly to *RBR* silencing, result in overproliferation. Contrary to the growth arrest upon RBR silencing, when exit from proliferation to differentiation is inhibited, the *e2fabc* mutant develops enlarged organs with supernumerary stem and differentiated cells as the quiescence is compromised. While E2F, RBR and the M-phase regulatory MYB3Rs are part of the DREAM repressor complexes, and recruited to overlapping groups of targets, they regulate distinct sets of genes. Only the loss of E2Fs but not the MYB3Rs interferes with quiescence, which might be due to the ability of E2Fs to control both G1-S and some key G2-M targets. We conclude that collectively the three canonical E2Fs in complex with RBR have central roles in establishing cellular quiescence during organ development, leading to enhanced plant growth.

## Introduction

In multicellular organisms, both the entry into and the exit from the cell division cycle is developmentally regulated. During plant development, there is a tight balance among cells proliferating in meristematic regions of emerging young organs, stem cells that are in a reversible quiescent status with a capacity to divide upon stimulation, and the most common cellular state of terminally differentiated cells formed after permanent exit from the cell cycle (Scheres, 2007). The principal regulation of cell proliferation by highly conserved molecular mechanisms is well established (Morgan, 2007), but how the cell cycle machinery is controlled in quiescent cells to maintain transient and stable repression of mitosis remains unclear.

Evidence from animal model organisms indicates that the RETINOBLASTOMA (Rb)-E2F transcriptional regulatory pathway plays a central role in coordinating proliferation and quiescence (Marescal & Cheeseman, 2020, Yao, 2014). Binding of Rb to the E2F-dimerization partner (DP) heterodimers can convert these complexes to transcriptional repressors, while phosphorylation of Rb by Cdk-cyclin complexes releases E2Fs and their ability to activate transcription. Accordingly, animal E2Fs can function as bistable switches on their target genes, determining whether the cells enter into the cell cycle or exit to establish quiescence (Yao et al., 2011).

The Rb-E2F pathway is remarkably conserved in plants, and it is well established that the plant RETINOBLASTOMA-RELATED (RBR) coordinates cell proliferation and differentiation by regulating the activity of three canonical E2F transcription factors (E2FA, E2FB and E2FC - (Borghi et al., 2010, Gutzat et al., 2011, Magyar et al., 2016, Wildwater et al., 2005) that form dimers with one of the two DP proteins (DPA and DPB). The three non-canonical E2Fs (E2FD, E and F) function independently of DP and RBR (Lammens al., 2009, Mariconti et al., 2002). Canonical E2Fs are structurally related to their animal counterparts, and consensus E2F-binding sites were identified in a large number of cell cycle regulatory genes (Mariconti et al., 2002, Vandepoele et al., 2002). Their C-termini contain the transactivation domain overlapping with the RBR-binding region (Kosugi & Ohashi, 2002, Magyar et al., 2000) and thus function as transcriptional activators or repressors depending on whether they are in complex with RBR (Magyar et al., 2012, Oszi et al., 2020). Overexpression studies suggest that E2FA and E2FB function as activators, but recent findings showed that they could also repress genes specifically involved in embryo and seed development (Leviczky et al., 2019). Surprisingly, genetic studies showed that E2F mutations either in single, double or even triple mutants did not universally compromise cell proliferation (Heyman et al., 2011, Leviczky et al., 2019, Oszi et al., 2020, Wang et al., 2014).

It has been established that loss of function *rbr* mutants are gametophytic lethal due to abnormalities during meiosis and gametophyte development (Ebel et al., 2004). Manipulation of RBR level during sporophyte development severely compromises the balance between proliferation and differentiation; ectopic overexpression of RBR led to diminishment of root and shoot meristems, while the consequence of RBR silencing is the inhibition of differentiation and increased number of stem cells (Wildwater et al., 2005, Wyrzykowska et al., 2006). Conditional RBR silencing strongly represses leaf growth by stimulating the overproliferation of stem cells in the stomatal lineage and inhibiting terminal differentiation (Borghi et al., 2010, Desvoyes et al., 2006), and triggers cell death (Horvath et al., 2017, Wildwater et al., 2005). It was suggested that upon RBR silencing the E2Fs are de-repressed, which is responsible for the above described developmental defects. However, the recent description of a fully viable, although partially sterile mutant plant, in which all three canonical E2Fs carry a T-DNA insertion (a triple *e2fabc* mutant) questions whether E2Fs are indeed essential for the activation of cell proliferation (Wang et al., 2014, Yao et al., 2018). This is surprising because there is ample evidence that plant E2Fs are the primary effectors of RBR just like in animals (Desvoyes & Gutierrez, 2020, Kent & Leone, 2019).

To understand the role of E2Fs for the developmental regulation of cell cycle, in this work we analysed the *e2fabc* triple mutant in detail for cell proliferation defects and for the deregulation of E2F targets. Surprisingly, just as upon RBR silencing, we observed overproliferation in the *e2fabc* triple mutant, suggesting that E2Fs are not required for maintaining cell proliferation, but rather have a repressive role. RBR silencing leads to growth arrest during seedling establishment due to the inhibition of this developmental transition and of cellular differentiation (Gutzat et al., 2011). This is not the case for the *e2fabc* mutant, which shows hyperplasia during embryonic and post-embryonic organ development with delayed cell cycle exit and unstable quiescence both in stem and in terminally differentiated cells. We established that the three E2Fs and RBR have largely overlapping gene targets and cell cycle genes were markedly upregulated in *e2fabc* mutant seedlings. We conclude that the three canonical E2Fs are collectively involved in cell cycle exit and in establishing transient and stable cellular quiescence by forming repressor complexes with RBR.

## Results

### The triple e2fabc mutant displays hyperplasia, leading to enlarged organs and plant stature

Several alleles of T-DNA insertion lines have been described for canonical E2Fs, all of which disrupt the carboxyl terminus containing the entire transactivation and RBR-binding domains (Extended Data Fig.1a, (Leviczky et al., 2019). As described previously, the triple *e2fabc* mutant is partially sterile, mostly due to aborted cell divisions in developing gametophytes (Yao et al., 2018), Fig.1a). In addition, at a very low frequency, we also observed supernumerary nuclei in the *e2fabc* mutant embryo sacs (2%; n=240), a characteristic phenotype for *rbr* mutant ovules Fig.1a; (Ebel et al., 2004, Zhao et al., 2017). Further similarities to *rbr* mutant phenotypes (Zhao et al., 2017) are the twin embryo sacs that may originate from two megagametophyte mother cells (Fig.1a; (Yao et al., 2018). These overproliferation phenotypes suggest that the canonical E2Fs may also have common repressor functions with RBR at some stages of gametophytic development.

**Figure 1:**
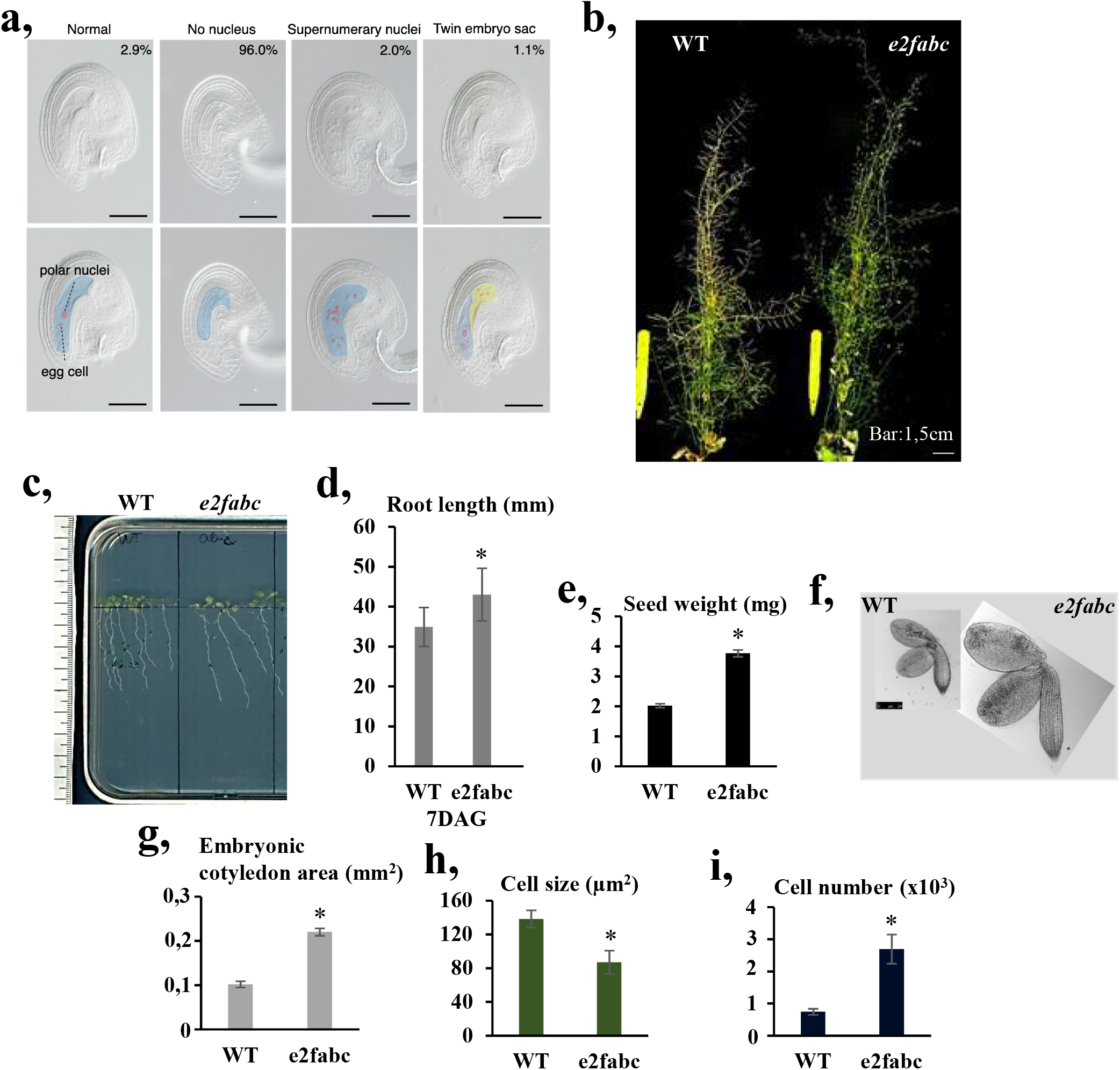
*E2fabc* mutants grow larger than the wild-type due to enhanced cell division. a: Gametophytic defects observed in *e2fabc* triple mutants (highlighted in the lower panel of pictures). DIC images of cleared ovules showing embryo sacs with non-distinguishable nuclei (96% of images), a minority of normal-looking embryo sacs that account for the few viable seeds produced by the mutant (2.9%), but also embryo sacs with multiple nuclei (2,0%) or even twin embryo sacs (1,1%), phenotypes that were also reported in *rbr* mutants. Scale bars = 50 μm. b: Pictures of mature plants showing that *e2fabc* mutants have shorter siliques but grow taller than the WT. Pictures were taken 30 DAG. c, d: Root length is increased in *e2fabc* mutants as shown by representative pictures of plantlets at 7 DAG (c), and quantification of root length (d), data are average +/− standard deviation (n=3 biological replicates; N=10 samples in each), \**P*≤0.05 indicates statistically relevant differences between the mutant and the WT (two-tailed paired *t*-test between the mutant and the WT). e: Seed weight is increased in *e2fabc* mutant, (weight of seeds is per 100 seeds, n=10/line). \*\**P*≤0.01 was considered significant between the WT and the mutant (two-tailed paired t-test between the mutant and the WT). f-i: *e2fabc* embryos are larger than those of WT and consist of more, smaller cells. f: representative pictures of dissected embryos from mature seeds, g: cotyledon area, h: epidermal cell size, i: epidermal cell number in embryonic cotyledon. For all graphs, data are average +/− standard deviation (n=3 biological replicates; N≥10 sample in each) and \**P*≤0.05, \*\**P*≤0.01, \*\*\**P*≤0.001 (two-tailed paired t-test between the mutant and the WT) indicate statistically relevant differences.

To our surprise, the *e2fabc* triple mutant was not only viable, but showed an increased plant size at the adult stage (Fig.1b) as well as longer roots (Fig.1c,d) and larger plantlet and rosette size (Extended Data Fig.2a-e). Importantly, another triple *e2fabc* mutant obtained by crossing the *e2fab* double with the *e2fc-2* T-DNA insertion mutant (Extended Data Fig.1), gave identical phenotypes with the original *e2fabc* mutant line (Extended Data Fig2f,g).

Due to the above described sterility, the *e2fabc* mutant only produced few seeds, but these were significantly larger and heavier than WT (Fig.1e; (Wang et al., 2014)), containing enlarged embryos (Fig.1f) that consisted of smaller and more numerous cells than WT (Fig.1g-i and Extended Data 2f-i-l). Likewise, developing leaves of the *e2fabc* mutant seedlings became gradually larger than those of WT, until they reached 1.5 times the size of WT leaves at 20 DAG (Fig.2a-b). Cellular analysis revealed that cell size was comparable with WT at early leaf development, but decreased at 16 DAG. We observed some fully differentiated lobbed pavement cells with straight division planes in the *e2fabc* mutant, but not in WT, indicating their re-entry into cell division (Fig.2e-f and Extended Data. Fig. 3a). The number of these extra divisions continuously rose and became the highest in the most advanced leaves at 16 DAG, resulting in smaller sized cells than in the WT (Fig.2f-d). As a result, the *e2fabc* mutant leaves contained almost twice as many cells (48%) than the control WT at 16 DAG (Fig.2c-d). Leaf cells dispersed among lobbed pavement cells, round and smaller than 60μm^2^ are considered to be stomatal meristemoids (Dong et al., 2009). The number of these cells was significantly increased in the expanding *e2fabc* mutant leaf at 12 and 16 DAG (Fig.2e-g), suggesting that not only the stable quiescence in pavement cells but also the transient quiescence in stomata stem cells is broken in the *e2fabc* mutant. Consistent with the overproliferation phenotypes, the expression levels of the S-phase-related *MCM3* (Extended Data Fig.3b) and the mitotic *CDKB1;1* (Fig.2h), continuously increased in the *e2fabc* mutant during leaf development whereas these were only detectable at the youngest stage of 9 DAG in the wild-type. The elevated expression of two genes associated with the stem cell lineage of stomatal meristemoids: *TOO MANY MOUTH (TMM;* Fig.2i), and *SPEECHLESS* (*SPCH*; Extended Data Fig.3b) was also in agreement with the observed phenotype. Taken together, these data suggest that E2Fs are essential to allow cell cycle arrest to establish both stable and transient quiescence.

**Figure 2:**
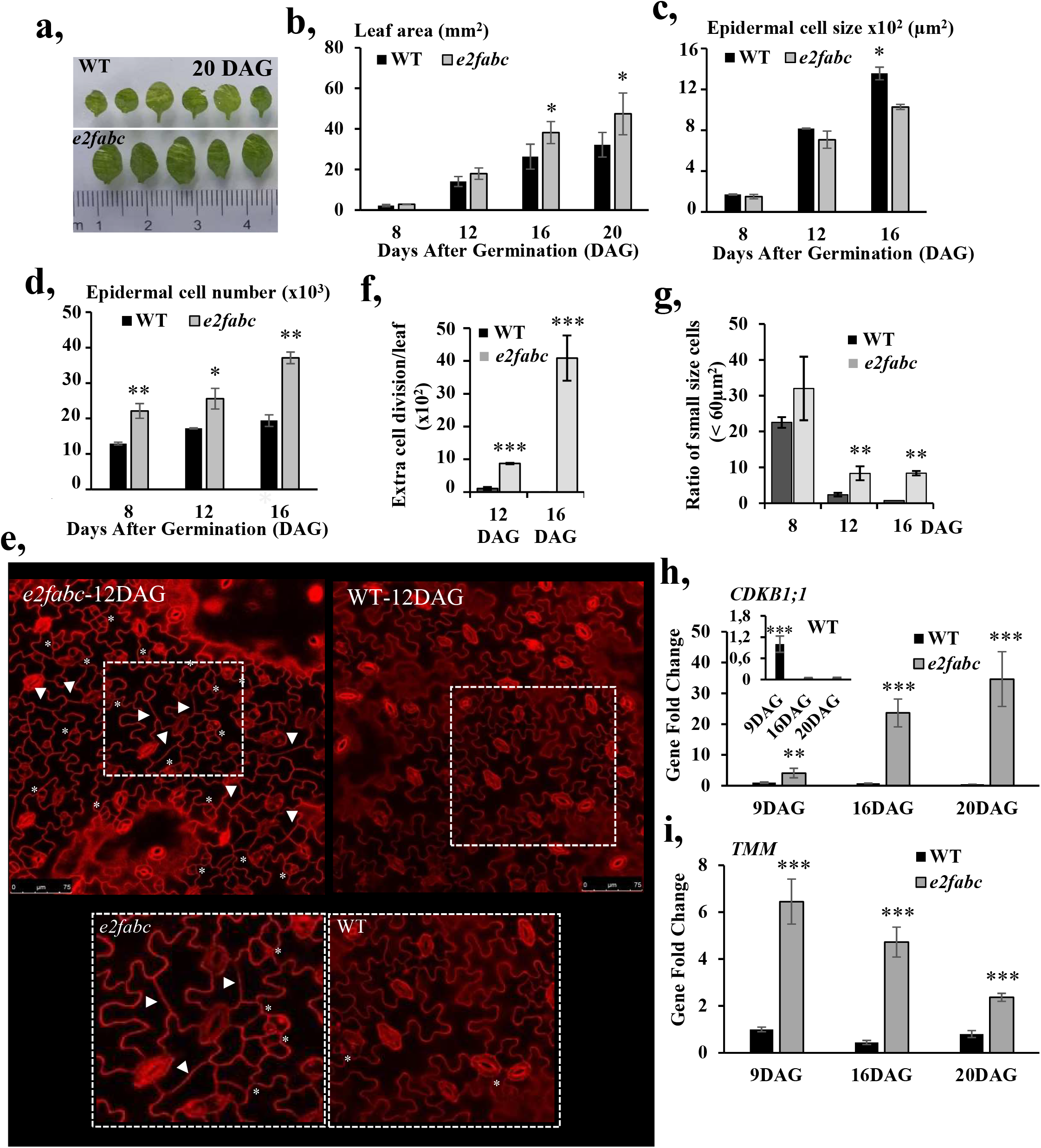
Cell proliferation is hyperactivated in developing leaves of *e2fabc* mutants. a-d: First leaves of *e2fabc* mutants are enlarged due to enhanced cell proliferation. a: representative pictures of the first leaves of WT plants or *e2fabc* mutants at 20 DAG. b: quantification of leaf area, c: quantification of cell area, and d: quantification of cell number in the leaf epidermis (first leaf pair). For all graphs, data are average +/− standard deviation n=3 biological replicates, N=10 samples in each. e-g: Cell division is reactivated in leaves of *e2fabc* mutants. e: Confocal microscope image of leaf epidermis of WT and *e2fabc* plants stained with propidium iodide (PI). Clustered stomata meristemoids (*) and division of differentiated puzzle-shaped cells (arrowheads) are observed in the mutant. f: Quantification of extra cell division events (n=3 biological replicates, N=6 samples in each), g: proportion of small cells corresponding to stomata meristemoids. Data are mean +/− sd., n=3 biological replicates, N=5 samples in each and sample size ≥600 cells \**P*<0.05, \*\**P*<0,01, \*\*\**P*<0,001 (two-tailed, paired t-test between the mutant and the WT at a given developmental time point). h-i: The cell cycle gene *CDKB1;1* and the stomata development gene *TMM* are over-expressed in *e2fabc* mutants compared to the wild-type. Expression of the two genes was monitored by qRT-PCR. Values represent fold changes normalised to the value of the relevant transcript of the wild type at 9DAG, which was set arbitrarily at 1. Data are means +/− sd., n=3 biological repeats. \*\**P*<0.01;, *** *P* < 0,001 (two-tailed, paired *t*-test between the WT and the mutant at a given time point). Abbreviations and primer sequences are listed in External Data Table 1.

### Canonical E2Fs maintain quiescence through the recruitment of RBR to repress cell cycle-related genes

To further understand the underlying mechanism for the repressor role of E2Fs, we investigated global gene expression changes in the triple *e2fabc* mutant seedlings. 1,767 genes were significantly upregulated and only 646 were downregulated in *e2fabc* compared to the WT (adj*P* value < 0.01). Additionally, Gene Ontology (GO) terms associated with upregulated, but not with downregulated genes, were highly enriched with function linked to cell cycle control (Fig.3a). To go further, we evaluated the expression changes of two gene categories: canonical E2F targets, defined as genes mis-regulated in *E2FA* over-expressing lines, and harbouring a consensus E2F binding motif in their promoter (Vandepoele et al., 2002), that are highly enriched in genes related to the control of the G1/S transition, and G2/M-specific genes, respectively (Haga et al., 2011). Among the genes upregulated in *e2fabc*, we found significant enrichment for canonical E2F target genes (*P*-value < 2.2e-16), but interestingly G2/M-specific genes were also affected (*P*-value = 0.0001401, Fig.3b). On the other hand, neither category was enriched among the downregulated genes in *e2fabc* mutant (Fig.3b, p = 0.3336 and 1 respectively). Similarly, scatter plot analysis comparing expression in *e2fabc* and wild type seedlings showed general upregulation of the canonical E2F target genes (Fig.3c).

**Figure 3:**
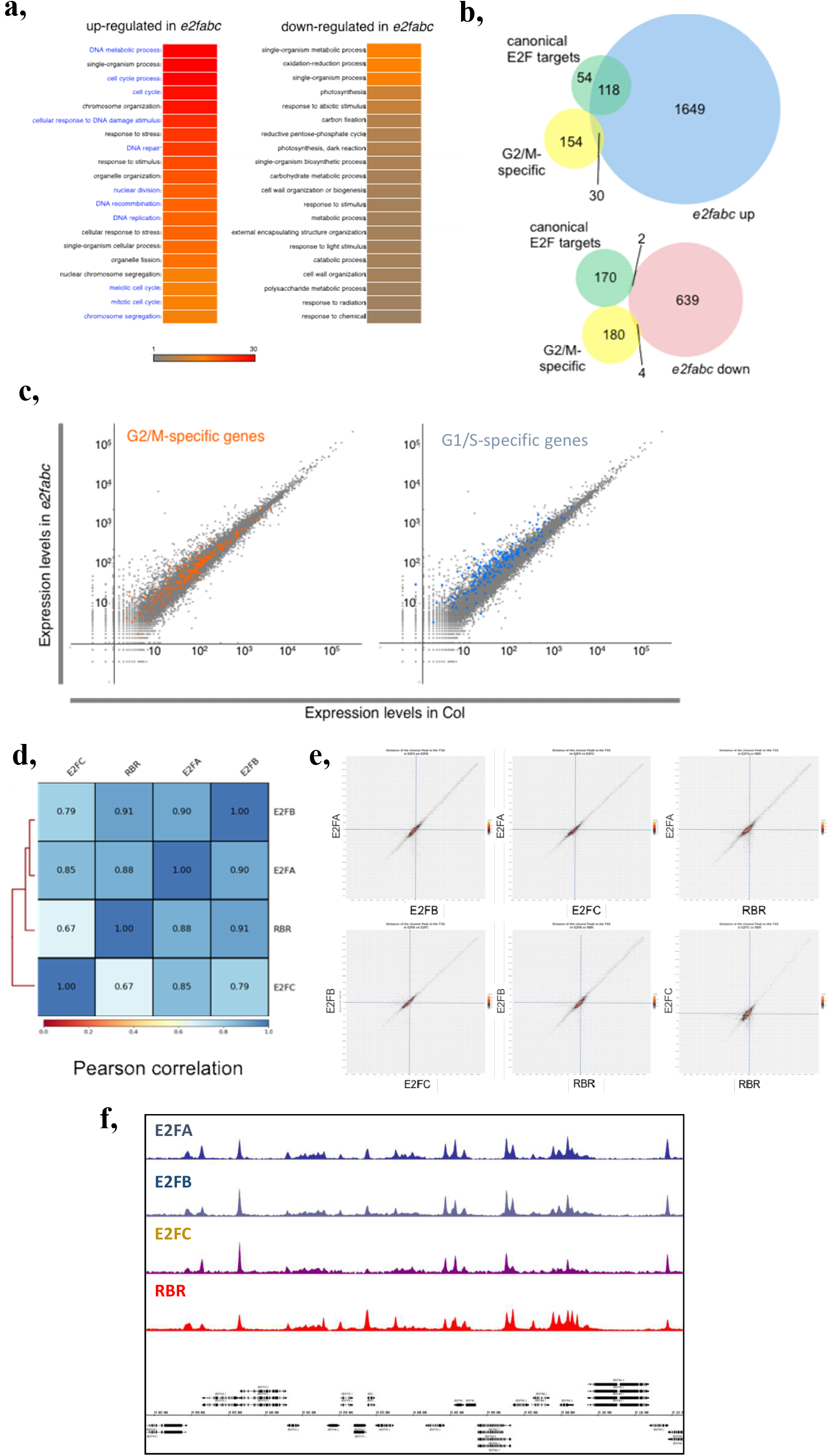
E2FA, B and C behave as repressors of cell cycle genes. a-c: Cell cycle genes are globally up-regulated in *e2fabc* mutants. a: GO term enrichment found amongst genes up or down-regulated in *e2fabc* triple mutants. The colour scale corresponds to the log_10_ of the FDR. b: Venn diagram showing the overlap between up or down-regulated genes in *e2abc* mutants and G1/S or G2/M cell cycle regulators. c: Scatterplot highlighting the expression of G2/M specific (orange) and canonical E2F target (blue) genes in *e2fabc* mutant compared to the WT. d-f: E2FA, B, C and RBR have largely overlapping target sites genome-wide. d: Pearson correlation of binding sites identified by ChIP-seq for E2FA, B, C and RBR. e: plots showing 2 by 2 comparison of the position of binding sites for E2FA, B, C and RBR on their targets. f: screenshot illustrating the similarity of ChIP-seq profiles obtained for E2FA, B, C and RBR.

Strong up-regulation of cell cycle genes is also observed when RBR function is inhibited (Borghi et al., 2010, Gutzat et al., 2011). To test whether E2FA, B, C and RBR actually target the same loci, we performed chromatin immunoprecipitation (ChIP) followed by sequencing (ChIP-seq) using plants expressing translational GFP fusions under the control of the native promoters (Kallai et al., 2020, Lang et al., 2021, Magyar et al., 2012, Oszi et al., 2020). Binding of all four proteins occurs mostly just upstream from transcriptional start sites of genes (Extended Data Fig.4a), and targets identified here for E2FA and RBR largely overlapped with previous reports (Verkest et al., 2014); Extended Data Fig.4b-c), validating the reliability of our data. Pearson correlation analysis (Fig.3d), as well as plots representing 2 by 2 comparisons of distances separating defined peaks from the closest TSS (Fig.3e) revealed that the four proteins have very similar binding profiles genome-wide, as further illustrated by the screenshot on Fig.3f.

The ability of canonical E2Fs and RBR to target the same set of genes, together with the observation that the simultaneous loss of E2FA, B and C triggers over-proliferation like RBR silencing, suggest that one key function of E2F proteins during vegetative development could be the recruitment of RBR to impose cell proliferation arrest. We therefore monitored the presence of RBR-DPB complexes in the *e2fabc* mutant, compared to *e2fab* mutants in which E2FC is still intact and functional. As expected, RBR was in complex with DPB in the *e2fab* double mutant (Fig.4a), but not in the *e2fabc* mutant, supporting that RBR recruitment to its targets is affected in these mutants.

**Figure 4:**
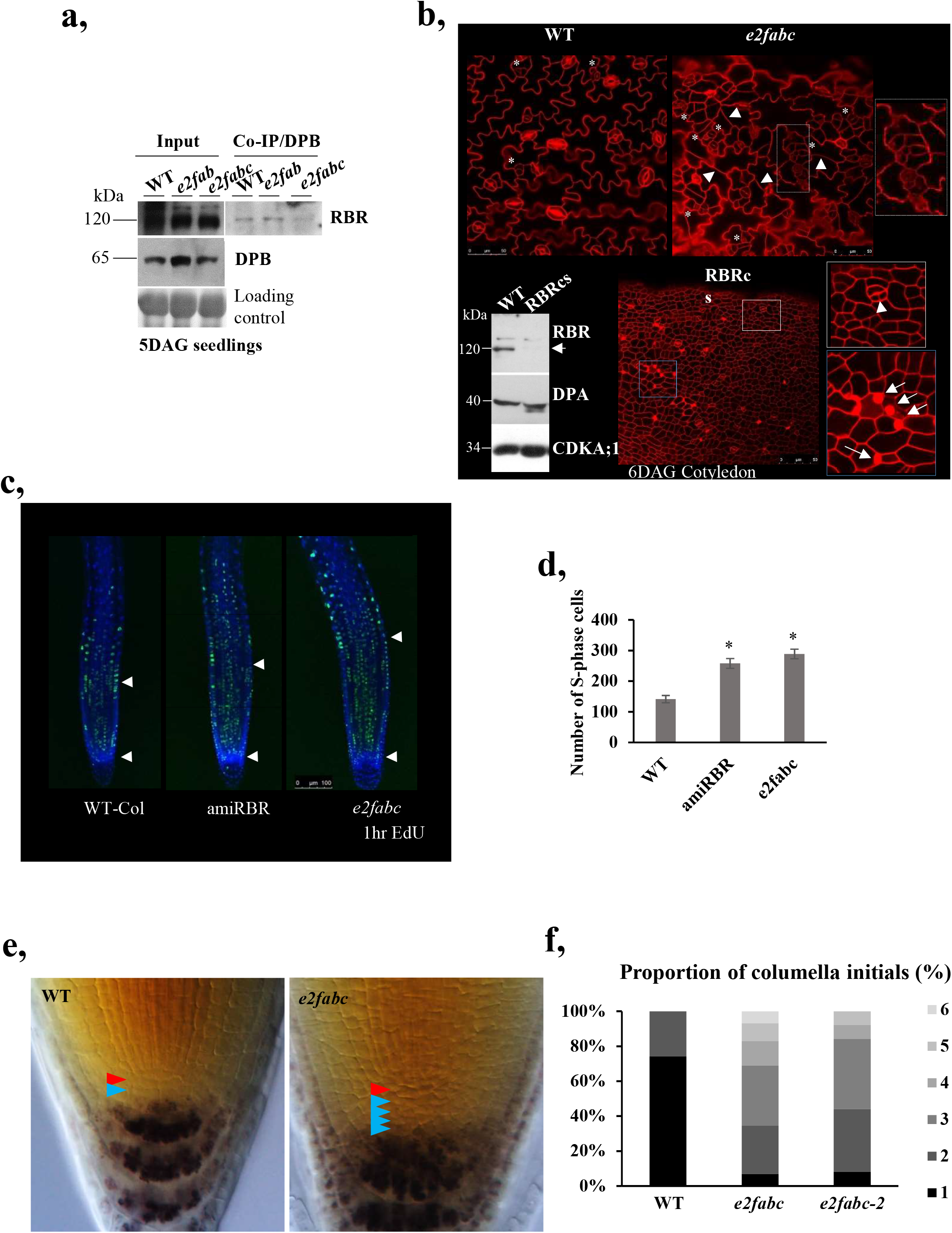
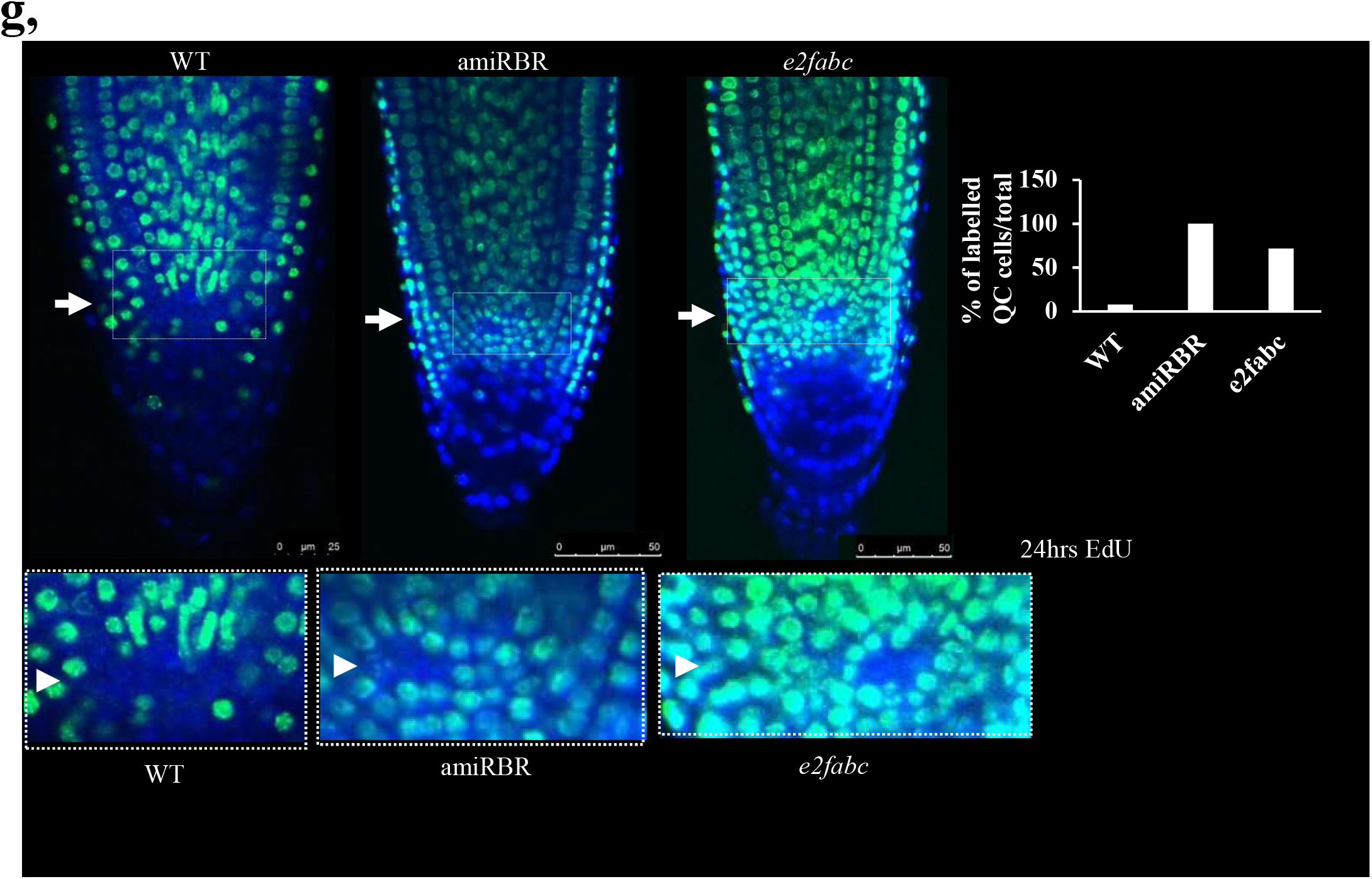
E2F/RBR complexes cooperate to control cellular quiescence. a: RBR-DPB complexes are almost undetectable in *e2fabc* mutants. Protein complexes were immuno-precipitated from protein extracts obtained from WT, *e2fab* or *e2fabc* mutant seedlings grown at 5 DAG using a native anti-DPB antibody. Comparable levels of both RBR and DPB can be detected in all genotypes (left, Input). However, although RBR could be co-IP-ed with DPB using protein extracts of WT or *e2fab* seedlings, amount of RBR immune-precipitated from *e2fabc* protein extracts were dramatically reduced. b: Comparison of the cellular phenotypes observed in the cotyledon epidermis of *e2fabc* triple mutants and *RBRcs* lines. Asterisks indicate clustered stomata meristemoids, arrowheads point to extra cell division events in puzzle-shaped differentiated pavement cells. In *RBRcs* lines all epidermal cells are small embryonic-like. c,d: *e2fabc* and *amiRBR* lines display enlarged root meristems. Roots of the indicated genotypes were labelled with EdU for 60 minutes. c: EdU staining of root tips of WT, *e2fabc* and amiRBR lines. White arrowheads mark the position of QC and the end of the meristematic zone indicated by enlarged nuclear size in the root epidermal cells. d: Graph showing the number of EdU labelled S-phase cells in these genetic backgrounds. Data are average +/− standard deviation (n=3 biological replicates, N=10 samples in each). * *P*< 0.05 (two-tailed, paired t-test between the mutant and the WT). e,f: *e2fabc* triple mutants accumulate supernumerary columella initials. e: microscopy images of lugol-stained root tips of WT and *e2fabc* mutants. Differentiated root cap cells can be identified thanks to amyloplast accumulation. The red arrowhead points to the QC and blue arrowheads indicate layers of columella initials. f: quantification of the proportion of roots with 1, 2, 3, 4, 5 or 6 layers of columella initials in WT, *e2fabc* or *e2fabc-2* mutants. g: Cell proliferation is reactivated in the QC of *e2fabc* and amiRBR lines. Plants were incubated in EdU for 24h. Representative images are shown on the top panel and the white arrow points at the QC. The graph below shows the proportion of roots showing EdU labelling in the QC (n = 100 samples in each).

Because the strong RBR co-suppression line is arrested during seedling growth *(RBRcs* - (Gutzat et al., 2011), we analysed the cotyledons for over-proliferation and compared them to the *e2fabc* mutant (Fig.4b, Extended Data Fig.5b-c). As in developing leaves, puzzle-formed pavement cells and guard cells in their regular bicellular forms could form in the cotyledon of *e2fabc* mutants, but meristemoids appeared clustered and differentiated pavement cells showed newly formed straight cross walls (Fig.4b, Extended Data Fig.5a). By contrast, most of the epidermal cells in *RBRcs* were small embryonic-like (Fig.4b; Extended Data Fig5d-e), and many dead cells could be visualised (Fig.4b; Extended Data Fig.5d-e; (Borghi et al., 2010, Gutzat et al., 2011). Cell size was markedly shifted from large cells towards smaller cells, and cell number was increased in both *RBRcs* and *e2fabc*, although much more in *RBRcs* (Extended Data Fig.6c-d). Consistently, the expression of key cell cycle regulators, DNA repair genes and embryonic genes was markedly increased in *RBRcs* lines, and to a lesser extent in *e2fabc* mutant (Extended Data Fig.7). Extra cell divisions could also be visualised both in cotyledons and young leaves of *e2fabc* triple mutant through the accumulation of the CYCB1;2 protein labelled with YFP (Extended Data Fig.8). By contrast, photosynthesis-related genes that fail to be activated in *RBRcs* line due to defects in cell differentiation, were expressed at comparable levels in *e2fabc* and WT plants (Extended Data Fig.7). These findings match the observation that cell differentiation is delayed but not compromised in *e2fabc* mutant whereas the *RBRcs* line is severely growth arrested because cell differentiation is prevented (Gutzat et al., 2011). Interestingly, late embryonic *LEC2* and *ABI3* genes were expressed at a comparable high level in both *RBRcs* and *e2fabc* mutants, and additionally seed storage 12S globulin was accumulated in the *e2fabc* seedlings suggesting that embryonic genes are repressed by RBR after germination probably in complex with E2Fs (Extended Data Fig.7b).

We also compared cell proliferation defects in the root meristem of *e2fabc* mutants and in a weaker RBR loss of function line, where RBR levels are reduced by an artificial microRNA (amiRBR, (Cruz-Ramirez et al., 2013)). Pulse labelling with 5-ethynyl 20-deoxyuridine (EdU) revealed a similar increase of S-phase cell number in the *e2fabc* mutant and the amiRBR line in comparison to the WT (Fig.4c-d). Furthermore, we observed supernumerary undifferentiated columella stem cells in the *e2fabc* root cap, as well as supernumerary cortex/endoderm initials (Fig.4e-f and Extended Data Fig.9a), as previously shown in an RBR silenced line (Wildwater et al., 2005). Mitosis, as followed by the YFP-tagged CYCB1;2 marker was clearly present in these supernumerary columella initials, providing further evidence for their sustained ability to divide (Extended Data Fig.9b). The lack of signal in quiescent centre (QC) compared to the surrounding cells when using prolonged EdU labelling allows to visualise the low frequency of S-phase in these cells, as shown in WT, while both in the *e2fabc* triple mutant and RBR silenced QC cells positive EdU labelled nuclei can readily be observed (Fig.4g). Quiescence of stem cells was shown to protect them against DNA damage (Cruz-Ramirez et al., 2013). As it was shown before for amiRBR, we found an increased sensitivity of root stem cells of the *e2fabc* line compared to WT to the DNA damaging agent of cisplatin (Extended Data Fig.10a-b). The RBR silenced lines also display spontaneous cell death within the root meristem, but this is not the case for the *e2fabc* line. Together, these data indicate that proliferation activities of the root meristem and specifically of the stem cells increased in the triple *e2fabc* mutant root, as is the case in amiRBR lines, providing evidence for the role of E2F/RBR complexes in the maintenance of the quiescent state.

### Though *E2Fs* and MYB3Rs can be in the same complex and recruited to a largely overlapping set of targets, they have unique regulatory roles

MYB3Rs together with E2FB, E2FC, and RBR, but not E2FA have been implicated in the repression of cell cycle genes as part of large multiprotein complexes, known as DREAM (Horvath et al., 2017, Kobayashi et al., 2015, Lang et al., 2021). We therefore investigated the interplay among E2Fs, RBR and repressive MYB3Rs in relation to their target gene regulation. We defined target genes as those detected with the highest confidence (*q*-value < 0.01 enrichment > 3). As expected, lists of genes bound by each E2F were largely overlapping, irrespective of the combination of isoforms looked at (Extended Data Fig. 11a). To collectively deal with target genes of E2FA, E2FB and E2FC, we defined a gene category representing general E2F targets, that is characterised by being amongst the most significant targets of at least two E2Fs (hereafter called E2F(2)). E2F(2)-, RBR- and MYB3R3-bound genes showed dramatic overlap in any combinations (Extended Data Fig.11b), defining 7 gene categories depending on the combination of factors that bind to them. *EMR* (bound by <u>E</u>2F, <u>M</u>YB3R3, and <u>R</u>BR) corresponded to 637 genes (Fig.5a). For most of these genes, peaks of ChIP signals exactly coincide for E2FA, E2FB, E2FC, RBR and MYB3R3 at the site of transcriptional initiation (Extended Data Fig.11c). GO terms associated with *EMR* genes were highly and exclusively enriched with categories related to cell cycle (Fig.5a). Enrichment of cell cycle-related GO terms was observed both for *EMR* and *EM*, and less prominently in *M*, but not at all in other categories (Fig.5a). Both the G1/S E2F targets and the G2/M-specific genes comprise a significant fraction of *EMR* and *EM* genes (Fig.5b). This suggests that genes with cell cycle-related functions are preferentially regulated by protein complexes containing both E2F and MYB3R3 transcription factors.

**Figure 5:**
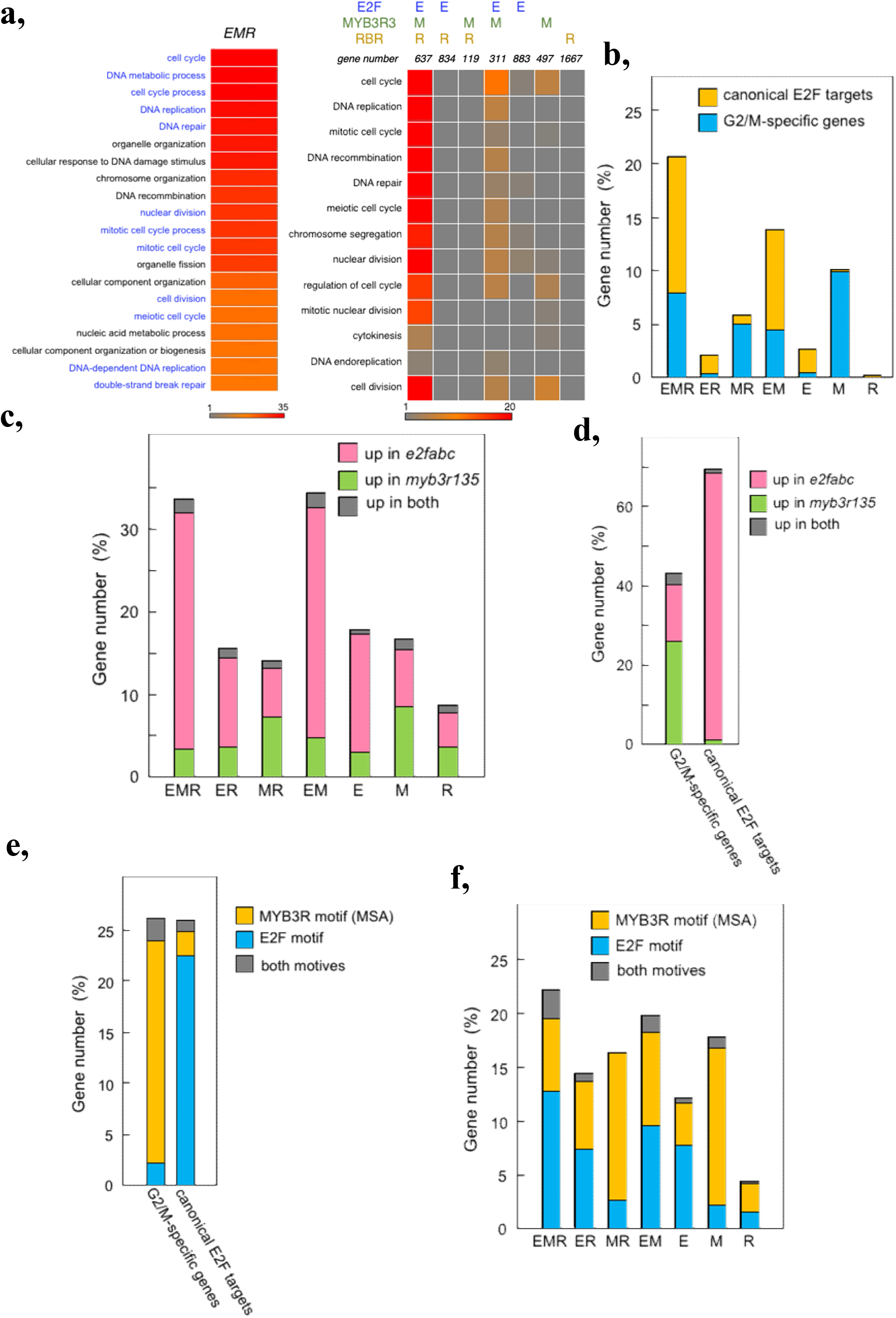
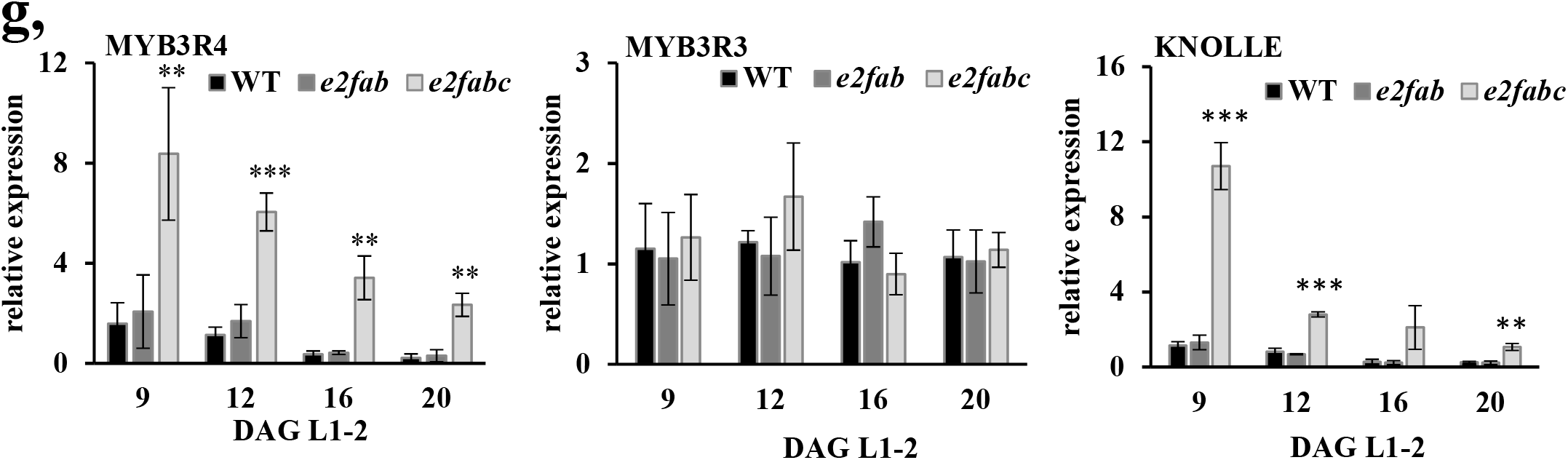
E2Fs and repressive MYB3Rs control distinct cell cycle genes, even though their target lists largely overlap. a: GO term enrichment analysis shows that genes bound by E2Fs, RBR, and MYB3Rs are significantly overrepresented for cell-cycle related genes, whereas this enrichment is not as obvious for genes targeted by 2 or only one of these factors. b: G1/S and G2/M-related genes are highly enriched those of the EMR category. This graph represents the percentage of G1/S (yellow), or G2/M (blue) genes amongst the six categories of genes defined according to their binding by E2F, RBR or MYB3Rs. c: Genes targeted by E2Fs, RBR and/or MYB3R are preferentially regulated either by E2Fs or by MYB3Rs. This graph represents the percentage of genes in each category that are up-regulated either in *e2fabc* or *myb3R1,3,5* mutants, or up-regulated in both. d,e,f: Cell-cycle related genes targeted by E2Fs, RBR and/or MYB3R are preferentially regulated either by E2Fs or by MYB3Rs. d: graph representing the percentage of G1/S or G2/M genes that are upregulated either in *e2fabc* or *myb3r1,3,5* mutants, or up-regulated in both. e: Graph representing the percentage of G2/M and G1/S genes harbouring a canonical E2F- or MYB3R-binding motifs in their promoters. f: Graph representing the percentage of genes harbouring a canonical E2F or MYB3R binding motif in their promoters within the 6 categories defined according to the binding profiles of E2Fs, RBR and MYB3Rs. g: Activating MYB3Rs but repressor MYB3Rs are up-regulated in *e2fabc* mutants. Expression of *MYB3R4, MYB3R3* and the MYB3R4 target and G2/M marker *KNOLLE* was monitored by qRT-PCR in the first leaf pairs of WT, *e2fab* and *e2fabc* mutants at 9, 12, 16 and 20 DAG. Values represent fold changes normalised to the value of the relevant transcript of the WT at 9DAG, which was set arbitrarily at 1. Data are means +/− sd., n=3 biological repeats. \*\**P*< 0.015;, *** *P*< 0,001 (two-tailed, paired *t*-test between the WT and the mutant at a given time point). Abbreviations and primer sequences are listed in External Data Table 1.

We then analysed the effects of E2F and MYB3R mutations on the expression of genes in each category as defined by our ChIP-seq data. For this purpose, RNA-seq gene expression datasets of *e2fabc* and *myb3r135* triple mutant were utilised. Genes within the *EMR* and *EM* groups were frequently upregulated in either *e2fabc* or *myb3r135* mutants, but very rarely in both (Fig.5c). Therefore, we concluded that in spite of the fact that MYB3Rs and E2Fs can be present in the same protein complex and they are recruited onto the same promoter site, on a particular target gene only one or the other regulate the expression. This unique mode of regulation can be also confirmed for canonical E2F target and G2/M-specific genes, which are preferentially influenced by E2Fs and MYB3Rs, respectively, but much less frequently by both factors (Fig.5d). Interestingly, the propension of genes to be influenced either by E2Fs or MYB3Rs correlated with the presence of consensus motives for E2F or MYB3R binding, respectively (Fig.5.e-f). Altogether, these data indicate that E2F and MYB3R transcription factors could bind to common cell cycle genes, but they specifically regulate the expressions of genes holding their characteristic E2F or MSA binding sequences, respectively.

While MYB3Rs are specifically targeted and regulate mitotic genes containing MSA elements, E2Fs can regulate cell-cycle genes involved both in G1-S and G2-M transitions (Fig.3b), an example is CDKB1:1 (Fig.2h), a critical controller of G2-M and the switch from mitosis to endoreduplication (Boudolf et al., 2004). Interestingly, the activating MYB3R, MYB3R4 is also a direct target of canonical E2Fs and is upregulated in the *e2fabc* mutant leaves at all analysed time points, whereas expression of the repressor MYB3R3 remained unchanged (Fig.5g). Accordingly, the MYB3R4 target, *KNOLLE* with a role in cytokinesis, was also upregulated (Fig.5g). These suggest that E2Fs may also regulate mitosis indirectly through the activating MYB3Rs. The ability of RBR and the three canonical E2Fs to regulate both G1/S and G2/M transitions may explain why only upon their loss the quiescence is broken, but not upon MYB3Rs that only act on mitotic genes.

## Discussion

The E2F-Rb regulatory pathway is a pivotal conserved regulator of cell proliferation both in animal and plant cells, acting at the critical transition point of G1 to S phase, known as restriction point. The textbook model is that E2Fs, when Rb-bound, restrict cell proliferation by preventing the expression of key cell cycle genes this is relieved through Rb phosphorylation by CDKs, the activity of which is regulated by a plethora of signalling inputs. Therefore, it is surprising that genetic knockout of all three canonical E2Fs in Arabidopsis can produce viable plants with a functional cell cycle. Here we show that while none of these E2Fs are required for activation of cell cycle genes, collectively they are responsible for their repression and thereby to establish both transient quiescence in stem cells as stable quiescence in differentiated cells. This scenario resembles to what has been established for the mammalian activator E2Fs (E2F1-3), as on one hand they were also found dispensable for cell proliferation while on the other they form complexes with Rb in differentiating cells to silence cell cycle genes (Chong et al., 2009).

Much of the previous studies have focused on the functional differences among the three canonical E2Fs (De Veylder et al., 2002, del Pozo et al., 2006, Magyar et al., 2005, Magyar et al., 2012, Sozzani et al., 2006). E2FA and E2FB are portrayed as transcriptional activators and E2FC as a repressor. Moreover, it appears that they work through different mechanisms, only E2FB and E2FC but not E2FA are part of DREAM-like multi-subunit protein complexes (Kobayashi et al., 2015, Lang et al., 2021). Cell proliferation in single *e2fa, e2fb* or *e2fc* mutants is largely normal and these plants only display mild phenotypes, e.g., the compromised meristem maintenance and reduced formation of lateral root primordia in *e2fa* mutant, and a slight increase in cell number during leaf development in *e2fb* mutant and an *e2fc* silenced line, (Berckmans et al., 2011, del Pozo et al., 2006, Leviczky et al., 2019, Oszi et al., 2020), suggesting that the distinction between activator and repressor classes of E2Fs is more nuanced. Interestingly, *e2fae2fb* double knock-out line could not be obtained (Li et al., 2017), but this appears to be allele specific, as combining the *e2fb* mutant with *e2fa-2* is viable and did not show proliferation defects during embryo development (Heyman et al., 2011, Leviczky et al., 2019). Here we show that combining this *e2fae2fb* double mutant with two different *e2fc* knockout lines, both display overproliferation, mostly due to the inability of maintaining quiescence. This suggests that all the three canonical E2Fs are required for repression of cell cycle genes even if they act through different molecular mechanisms (Leviczky et al., 2019, Oszi et al., 2020). The principal component of repressor complexes on all canonical E2Fs is RBR and it was demonstrated that all three canonical Arabidopsis E2Fs strongly interact with it (Lang et al., 2021, Magyar et al., 2012, Oszi et al., 2020). However, we have seen a stunning difference between the phenotypes of RBR silencing and the triple *e2fabc* mutant, both showing overproliferation during organ development, but only upon RBR silencing and not in the *e2fabc* mutant is cell differentiation repressed. A similar inhibition of differentiation occurs when CYCD3;1, the regulatory component of the major RBR kinase is overexpressed (Dewitte et al., 2003). On the one hand, RBR repression on E2Fs is the pivotal regulator of the restriction point in G1 and in response to developmental cues can determine the G1 length and whether cells can exit to differentiation. In this scenario tuning E2F activation of cell cycle genes by RBR is the critical mechanism. On the other hand, mutation of all three E2Fs abrogates both the RBR-E2F corepressor function and the transcriptional activation of cell cycle genes. The comparison of RBR silencing and *e2fabc* mutant lines indicates that E2F activator function is important for the cell cycle exit and onset of differentiation, while maintaining quiescence is primarily regulated by RBR-E2F co-repression. In addition, our results show that plant E2Fs, like their animal counterparts, are the primary effectors of RBR in nearly every process where RBR is involved (Fig.6a). The transient stem cell quiescence is most sensitively abrogated in the stomata lineage when RBR is silenced or the canonical E2Fs are mutated, as indicated by the abundance of small cells characterised as stomata stem cells and by the continuous upregulation of stomata stem cell factors, *TMM* and *SPCH* (Borghi et al., 2010). Interestingly, RBR, in an E2F-independent manner, by direct interaction with the stomata terminal differentiation transcription factor, FAMA, was also shown to regulate the late steps of differentiation within the stomata cell linage (Matos et al., 2014). However, in the *e2fabc* mutant we have not seen the characteristic phenotype of the *fama-1* mutant, called fama-tumours, clusters of small, narrow epidermal cells that are unable to go through the final stages of stomata differentiation. RBR and E2Fs can also directly regulate genes involved in late embryonic differentiation programs, e.g., *LEC2* and *ABI3*, which are direct E2F and RBR targets and are upregulated both when RBR is silenced or in the *e2fabc* mutant, but only in the former the developmental transition of seedling establishment is blocked (Gutzat et al., 2011). Therefore, it is likely that as during leaf emergence, the developmental arrest in seedlings, when RBR is silenced, is due to the deregulated cell cycle.

**Figure 6:**
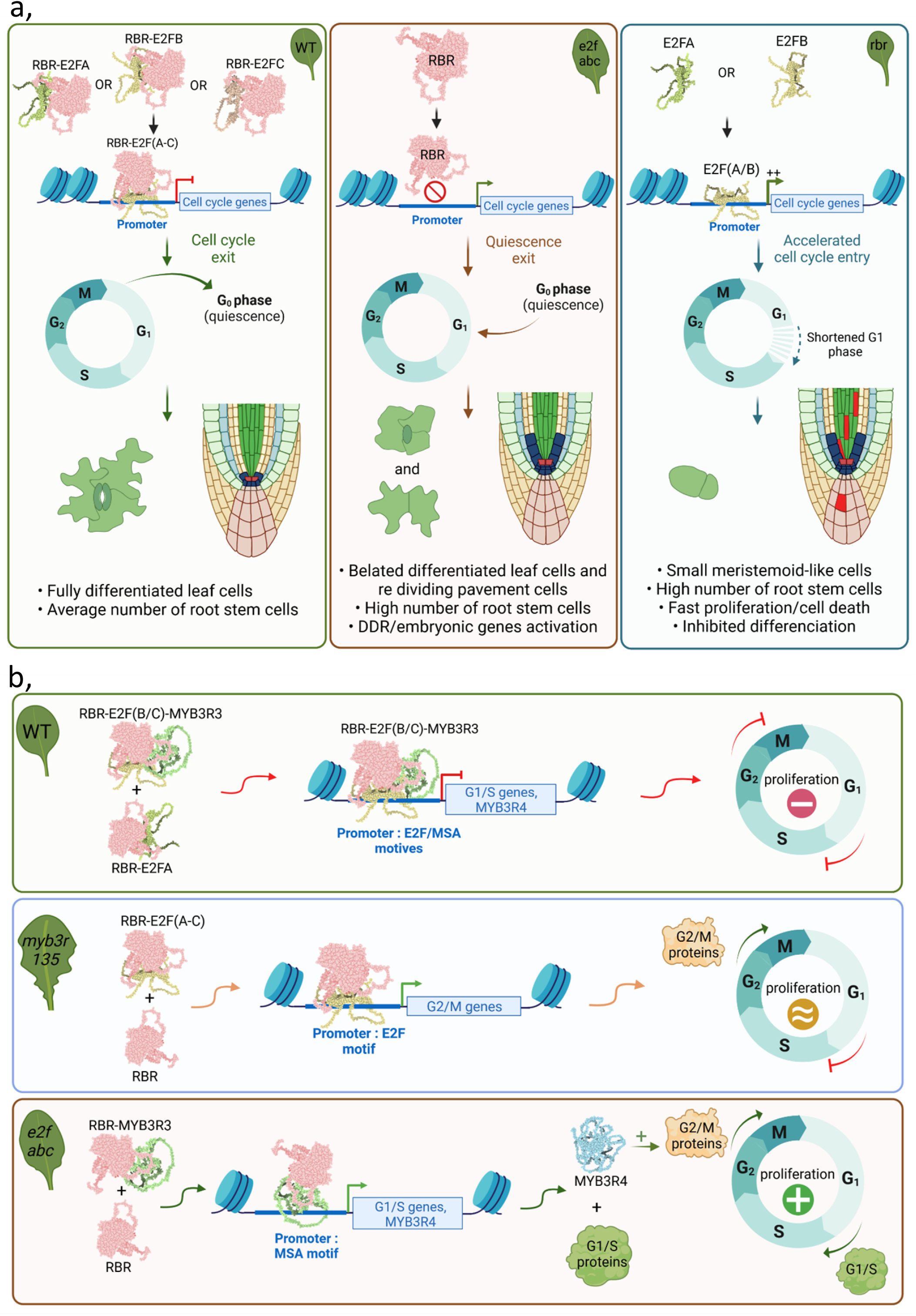
Schematic model how E2Fs and RBR induce developmental quiescence. a: Canonical E2Fs function as co-repressors of cell cycle genes with RBR. Left: in the wild-type, E2Fs form complexes with RBR to repress cell cycle genes and allow cell division arrest in quiescent and differentiated cells. Middle: in the e2fabc mutant, lack of RBR-E2F complexes results in de-repression of cell cycle genes inducing a delay in cell differentiation as well as division reactivation in quiescent cells. Right: in RBR-deficient lines, E2F/RBR complexes are also lacking, but E2Fs are fully active, leading to a more pronounced hyperactivation of cell cycle genes and thus completely preventing cell differentiation, and inducing cell death. b: Distinct molecular roles of E2Fs and MYB3Rs in the control of cell cycle arrest. Top: in the wild-type, Repressor complexes containing RBR, E2Fs and MYB3Rs target G1/S and G2/M genes to repress their expression either via the MSA or the E2F motives. Middle: in the *myb3R1,3,5* mutant that lacks repressor MYB3Rs, E2Fs are still present and able to repress G1/S and some G2/M genes bound via the E2F motif. Cell proliferation is therefore not induced in differentiated cells. Bottom: in the e2fabc mutant, repressor MYB3Rs can still repress some G2/M genes with or without RBR, but repression of both G1/S and G2/M activators such as MYB3R4 is lost, resulting in enhanced cell proliferation.

It was shown both in animals and plants that quiescence protects long-lived stem cells against stress and toxicities (Cheung & Rando, 2013, Cruz-Ramirez et al., 2013, Legesse-Miller et al., 2012). In agreement, we found that stem cells are sensitive to genotoxic stress both upon RBR silencing and in the *e2fabc* mutant. RBR and E2Fs were also implicated to directly regulate the expression of DNA damage genes and cell death. In this respect we observed a much more pronounced spontaneous cell death with RBR silenced cells than in the *e2fabc* mutant, which correlated with the level of induction of DNA damage genes, suggesting that both co-repression and E2F transactivation function contributes to this regulation.

RBR, E2FB and E2FC together with the mitotic MYB3Rs can be part of large repressor complexes, called DREAM (Lang et al., 2021). Here we show that RBR, E2Fs and MYB3Rs are also recruited to largely overlapping sets of cell cycle target genes, but still maintain a specific regulatory function on E2F targets or mitotic genes dependent on whether the MSA or E2F target sequence is present in these promoters. We have shown previously that mutation of the three repressor MYB3R (*myb3r1/3/5*) in Arabidopsis resulted in excess cells in developing organs but exclusively within the meristems (Kobayashi et al., 2015). Interestingly, mitotic genes like *CYCB1;2* were found to be ectopically expressed also in differentiated leaf cells, but this did not result in mitotic reactivation, possibly because E2F-RBR complexes are still present, and capable of repressing genes involved in the G1/S transition, which may explain why quiescence is efficiently maintained in the myb3r1/3/5 mutant (Figure 6b). In contrast, in the *e2fabc* mutant, where the E2F target genes are de-repressed, we find that both the transient quiescence in stem cells and terminal quiescence in differentiated cells are broken. Although E2Fs primarily regulate genes with G1-S function, they were also shown to control the expression of key G2-M regulatory genes, like the plant specific CDKBs, supporting that E2Fs regulate both cell cycle transitions (Magyar et al., 2005). Moreover, E2Fs and RBR were specifically enriched on the promoter of the activator MYB3R4, a key regulator of mitosis, and indeed it was induced and activated in the *e2fabc* mutant leaf, which might explain the central role of E2Fs to maintain cellular quiescence by controlling both G1/S and G2/M transitions (Figure 6b).

Taken together, we have shown that the canonical E2Fs together with RBR form a repressor complex on a cohort of cell cycle genes that is pivotal for maintaining quiescence. Through this repression mechanism it is possible to influence plant growth which could open novel ways for improving crop biomass and productivity.

## Methods

### Plant material and growth conditions

Arabidopsis thaliana Col-0 ecotype was the WT and background of all transgenic lines used in this study. In vitro-cultured plants were grown on a half-strength germination medium under continuous light at 22°C. Soil grown plants were cultivated in a greenhouse at 22°C under long-day conditions (16h light/8h dark). Most of the transgenic and T-DNA insertion mutant lines used in this study have been previously published: E2FA-GFP, RBR-GFP (Magyar et al., 2012), E2FB-GFP (Oszi et al., 2020), E2FC-GFP (Kallai et al., 2020), *e2fb-1* (SALK_103138), *e2fa-2* (GABI-348E09), *e2fc-1*, (GK-718E12,(Berckmans et al., 2011, Wang et al., 2014); the double *e2fab (e2fa-2/e2fb-1*) was reported by (Heyman et al., 2011), and the triple *e2fabc (e2fa-2/e2fb-1/e2fc-1*) was described in (Wang et al., 2014). The second T-DNA insertion line for E2FC, named as *e2fc-2* (SAIL-1216G10) was obtained from the SALK Institute and the *e2fabc-2* triple mutant was generated by crossing the *e2fab* double (*e2fa-2/e2fb-1*) with the *e2fc-2*. RBR cosuppression (*RBRcs*) seedlings were identified in transgenic lines expressing either the RBR-3xCFP (Leviczky et al., 2019) or the RBR-RFP (Biedermann et al., 2017) in the WT background under the control of the native RBR promoter showing strong growth arrested phenotype identical with previously reported (Gutzat et al., 2011).

Young seedlings (5, 7, or 14 days after germination, DAG) or the first true leaf pairs of the wild-type and mutant Arabidopsis lines (*e2fabc* and *e2fab*) grown *in vitro* were harvested 9, 12, 16 and 20 DAG, flash frozen and stored at −80°C.

### Microscopy

Mature dried seeds were imbibed for 1 h and dissected under the stereo-microscope. Isolated embryos were stained with PI and photographed under confocal laser microscopy (SP5, Leica). Organ and epidermal cell sizes were measured using ImageJ software (Leviczky et al., 2019, Schneider et al., 2012). Cotyledons and the first true leaf pairs of wild-type and mutant lines were dissected from seedlings at 6 DAG and 8-16 DAG, respectively. Leaves were stained with propidium iodide (PI; 20 μg/mL) and images on the abaxial side of the cotyledon or the leaf were taken and analysed by confocal laser microscopy. 600-300 cells were counted and measured per cotyledon or leaf samples (n=3 biological replicates and N=5 samples in each case were studied for the transgenic lines and the control WT) using ImageJ software. Average cell size was calculated, and the total cell number was extrapolated to the whole cotyledon and the first real leaf pair according to previously described methods (Asl et al., 2011, Oszi et al., 2020). Roots was also visualised after PI staining (20 μg/mL) under confocal laser microscopy (Oszi et al., 2020).

For DIC microscopy of cleared tissues, plant tissues were fixed in a 9:1 of ethanol and acetic acid solution and cleared with Hoyer’s solution (a mixture of 100 g chloral hydrate, 10 g glycerol, 15 g gum arabic, and 25 mL water). They were mounted onto glass slides and used for microscopic observations with differential interference contrast (DIC) microscope (BX51, Olympus). Images were captured with a CCD camera (DP74, Olympus) and an image capture software (CellSens Standard, Olympus). For visualization of starch granules in columella cells, roots were incubated for 3 min in Lugol solution (Sigma) before clearing.

For EdU incorporation assay WT and the mutant seedlings were grown in half strength liquid MS containing 10 μM EdU (Click-iT Alexa Fluor 647 Imaging Kit; Invitrogene) and incubated either for an hour or for 24 hours. Afterward, seedlings were treated according to (Vanstraelen et al., 2009), and the root samples were also stained by DAPI solution and the observations were done under the confocal laser microscope.

### cDNA preparation and RT-qPCR

RNA was extracted from the plant material using a CTAB-LiCl method of (Jaakola et al., 2001). RNA samples were treated with DNase1 (ThermoScientific #EN0521) according to the manufacturer’s protocol. cDNA was synthesised using 1 μg of RNA using the ThermoScientific Reverse Transcription Kit (#K1691) with random hexamers based on the manufacturer’s prescription. Mock reaction without RevertAid enzyme was also prepared to ensure that there is no contaminating genomic DNA in the samples. (RT-qPCR) in the presence of SYBR Green (TaKaRa TB Green Primer Ex TaqII, #RR820Q) was carried out using a BioScript PCR kit (Bioline) according to the manufacturer’s instructions in a BioRAD CFX 384 Thermal Cycler (BioRAD) with the following setup: 500C 2 min, 95°C 10 min, 95°C 15 sec, 60°C 1 min, 40 cycles followed by melting point analysis. Each reaction was carried out in tree technical replicates and reaction specificity was confirmed by the presence of a single peak in the melting curve. All the data were normalised to the average Ct value of two housekeeping genes (ACTIN and UBIQUITIN) and the calculated efficiency was added to the analysis. Amplification efficiencies were derived from the slope of amplification curves at the exponential phase. Primer sequences are summarised in Extended Data Table 1.

### Immunoprecipitation (IP) and immunoblotting

Total proteins were extracted from young seedlings and IP and immunoblotting assays were carried out as described (Oszi et al., 2020). Primary antibodies used in immunoblotting experiments were chicken anti-RBR antibody (1:2,000 dilution; Agrisera), mouse monoclonal anti-PSTAIRE (1:40,000 dilution, CDKA;1 specific; Sigma), anti-phospho-specific Rb (Ser-807/811) rabbit polyclonal antibody (1:500 dilution; Cell Signaling Tech), anti-DPA, anti-E2FB polyclonal rabbit antibodies (both 1:400 dilution, Magyar et al., 2005), anti-DPB polyclonal rabbit antibody (Umbrasaite et al., 2010), and anti-E2FA rat polyclonal antibody (1:400 dilution, (Leviczky et al., 2019), rabbit polyclonal antibody anti-12S globulin (1:10000; (Shimada et al., 2003). After the primary antibody reaction, the membrane was washed (TBST) and incubated with the appropriate secondary antibody conjugated with horseradish peroxidase at room temperature. Afterward, chemiluminescence substrate was applied according to the manufacturer description (SuperSignal West Pico Plus, Thermo Fisher Scientific) or Immobilon western horseradish peroxidase (Millipore). For IP, equal amounts of protein samples (800 μg) in the extraction buffer (Oszi et al., 2020) were incubated with anti-DPB antibodies for 1h at 4°C on a rotary shaker. Dynabeads Protein A (Invitrogen) was used to pull down polyclonal antibodies, and after washing the beads proteins were eluted by adding SDS sample buffer followed by 5 min boiling. Eluted proteins were loaded on SDS-PAGE gels (10%) and after protein gel electrophoresis they were immunoblotted as described (Oszi et al., 2020).

### Chromatin immunoprecipitation followed by high-throughput sequencing (ChIP-seq) assay

ChIP-seq assays were performed on 14-d-old seedlings using anti-GFP antibody (Abcam). Plant material was cross-linked in 1% (v/v) formaldehyde for 15 min under vacuum at room temperature and cross-linking was then quenched with 0.125 M glycine for 5 min. Cross-linked plantlets were ground in liquid nitrogen, and nuclei were lysed in Nuclei Lysis Buffer (0.1% SDS, 50 mm Tris-HCl at pH 8, 10 mm EDTA, pH 8). Chromatin was then sonicated for 5 min using a Covaris S220 (Peak Power: 175, cycles/burst: 200. Duty Factory: 20). Immunoprecipitation was performed overnight at 4°C with gentle shaking, and immunocomplexes were next incubated for 1h at 4°C with 40 μL of Dynabeads Protein A (Thermo Fisher Scientific). The beads were washed 2 × 5 min in ChIP Wash Buffer 1 (0.1% SDS, 1% Triton X-100, 20 mM Tris–HCl at pH 8, 2 mM EDTA at pH 8, 150 mM NaCl), 2 × 5 min in ChIP Wash Buffer 2 (0.1% SDS, 1% Triton X-100, 20 mM Tris–HCl at pH 8, 2 mM EDTA at pH 8, 500 mM NaCl), 2 × 5 min in ChIP Wash Buffer 3 (0.25 mM LiCl, 1% NP-40, 1% sodium deoxycholate, 10 mM Tris–HCl at pH 8, 1 mM EDTA at pH 8) and twice in TE (10 mm Tris-HCl at pH 8, 1 mM EDTA at pH 8). ChIPed DNA was eluted by two 15-min incubations at 65°C with 250 μL of Elution Buffer (1% SDS, 0.1 m NaHCO_3_). Chromatin was reverse crosslinked by adding 20 μL of NaCl 5 M and incubated overnight at 65°C. Reverse cross-linked DNA was treated with RNase and Proteinase K and extracted with phenol–chloroform. DNA was ethanol precipitated in the presence of 20 μg of glycogen and resuspended in 10 μL of nuclease-free water in a DNA low-bind tube. Libraries were then generated using 10 ng of DNA and NEBNext Ultra II DNA Library Prep Kit for Illumina (NEB), following the manufacturer’s instructions. The quality of libraries was assessed with an Agilent 2100 Bioanalyzer (Agilent), prior to 1 × 75 bp high-throughput sequencing by NextSeq 500 (Illumina).

### Analysis of ChIP-seq data

Trimmomatic-0.38 was used for quality trimming (Martins et al., 2011). Parameters for read quality filtering were set as follows: minimum length of 36 bp; mean Phred quality score greater than 30; leading and trailing bases removal with base quality <5. The reads were mapped onto the TAIR10 assembly using Bowtie 2 (Langmead & Salzberg, 2012) with mismatch permission of 1 bp. To identify significantly enriched regions, we used MACS2 (Gaspar, 2018). Parameters for peaks detection were set as follows: number of duplicate reads at a location:1; mfold of 5:50; Q-value cutoff: 0.05; extsize 200; sharp peak. Average scores across genomic regions were extracted using the multiBigwigSummary command of the DeepTools package after data normalization with the S3norm software (Xiang et al., 2020). Data were visualized using WashU and the plot Heatmap tool of the DeepTools package (Ramirez et al., 2016) was used to generate heatmaps while the ggplot2 package was used to draw metaplots.

### Motif enrichment analysis

Position weight matrix (PWM) data for binding motifs of E2FA and MYB3R5, determined by DNA affinity purification sequencing (DAP-Seq), were obtained from a website of Plant Cistrome Database at http://neomorph.salk.edu/PlantCistromeDB (O’Malley et al., 2016), and used as E2F- and MYB3R-binding motifs. Motif search based on PWM was conducted using the matchPWM function implementing in R Biostrings package. Forward and reverse of each motif were used as PWM with minimum score of 50%.

### Transcriptome analysis

Total RNAs were extracted using the RNeasy Plant Mini Kit (QIAGEN, Germany) from whole seedlings according to the manufacturer’s instructions. Total RNAs from WT and *e2fabc* at 9 DAG were used for construction of cDNA libraries using the TruSeq RNA Library Preparation Kit v2 (Illumina, United States) according to the manufacturer’s protocol. For transcriptome profiling in WT and *e2fabc* plants, we analyzed three biological replicates for statistical analysis. The libraries were sequenced using the NextSeq500 sequencer (Illumina, United States). Raw reads containing adapter sequences were trimmed using bcl2fastq (Illumina, United States), and nucleotides with low-quality (QV < 25) were masked by N using the original script. Reads shorter than 50 bp were discarded, and the remaining reads were mapped to the cDNA reference using Bowtie with the following parameters: “–all–best–strata” (Langmead et al., 2009). The reads were counted by transcript models. Differentially expressed genes were selected based on the adjusted P-value calculated using edgeR (version 3.20.9) with default settings (Robinson et al., 2010).

### Other methods

Gene ontology analysis was carried out using the singular enrichment analysis tool offered by agriGO with the default settings (Du et al., 2010).

## Data availability

RNA-Seq data for WT and *e2fabc* can be accessed from the DDBJ database under accession number DRA015182. ChIP-Seq data of RBR, E2FA, E2FB and E2FC can be accessed at Gene Expression Omnibus database under accession number GSE218481. Arabidopsis mutants and transgenic lines, as well as plasmids generated in this study are available from the corresponding author upon reasonable request.

## Acknowledgements

We thank Xinnian Dong for the *e2fabc* mutant line, Arp Schnittger for the RBR-RFP line.

This work was supported by a grant from the Hungarian National Research Funding (NKFI-139202) to Z. Magyar, the Japan Society for the Promotion of Science KAKENHI (22K06261 and 22H04714) to M. Ito, Japan Science and Technology Agency (JST grant number JPMJPF2102) to M. Ito, Biotechnology and Biological Sciences Research Council BBSRC-NSF grant BB/M025047/1 to L. Bogre and C Papdi.

## Author contributions

Z.M., L.B., M.I., C.R., and M.B. conceived the study; M.G., C.R, ZM., MI designed the experiments; MG., CR., ZM., YN., EM., RB-C., HT., AZ., DB., DL., KM., FN., HI., EŐ., and CP carried out the experiments; XH., JA. and TS analyzed the data, CB did the model; Z.M., C.R., M.G., L.B., M.I. and M.B. wrote the paper.

## Additional information

### Supplementary information

The online version contains supplementary material.

## References

Asl LK, Dhondt S, Boudolf V, Beemster GT, Beeckman T, Inze D, Govaerts W, De Veylder L (2011) Modelbased analysis of Arabidopsis leaf epidermal cells reveals distinct division and expansion patterns for pavement and guard cells. Plant Physiol 156: 2172–83

Berckmans B, Vassileva V, Schmid SP, Maes S, Parizot B, Naramoto S, Magyar Z, Alvim Kamei CL, Koncz C, Bogre L, Persiau G, De Jaeger G, Friml J, Simon R, Beeckman T, De Veylder L (2011) Auxin-dependent cell cycle reactivation through transcriptional regulation of Arabidopsis E2Fa by lateral organ boundary proteins. Plant Cell 23: 3671–83

Biedermann S, Harashima H, Chen P, Heese M, Bouyer D, Sofroni K, Schnittger A (2017) The retinoblastoma homolog RBR1 mediates localization of the repair protein RAD51 to DNA lesions in Arabidopsis. EMBO J 36: 1279–1297

Borghi L, Gutzat R, Futterer J, Laizet Y, Hennig L, Gruissem W (2010) Arabidopsis RETINOBLASTOMA-RELATED is required for stem cell maintenance, cell differentiation, and lateral organ production. Plant Cell 22: 1792–811

Boudolf V, Vlieghe K, Beemster GT, Magyar Z, Torres Acosta JA, Maes S, Van Der Schueren E, Inze D, De Veylder L (2004) The plant-specific cyclin-dependent kinase CDKB1;1 and transcription factor E2Fa-DPa control the balance of mitotically dividing and endoreduplicating cells in Arabidopsis. Plant Cell 16: 2683–92

Bouyer D, Heese M, Chen P, Harashima H, Roudier F, Gruttner C, Schnittger A (2018) Genome-wide identification of RETINOBLASTOMA RELATED 1 binding sites in Arabidopsis reveals novel DNA damage regulators. PLoS Genet 14: e1007797

Cheung TH, Rando TA (2013) Molecular regulation of stem cell quiescence. Nat Rev Mol Cell Biol 14: 329–40

Chong JL, Wenzel PL, Saenz-Robles MT, Nair V, Ferrey A, Hagan JP, Gomez YM, Sharma N, Chen HZ, Ouseph M, Wang SH, Trikha P, Culp B, Mezache L, Winton DJ, Sansom OJ, Chen D, Bremner R, Cantalupo PG, Robinson ML et al. (2009) E2f1-3 switch from activators in progenitor cells to repressors in differentiating cells. Nature 462: 930–4

Cruz-Ramirez A, Diaz-Trivino S, Wachsman G, Du Y, Arteaga-Vazquez M, Zhang H, Benjamins R, Blilou I, Neef AB, Chandler V, Scheres B (2013) A SCARECROW-RETINOBLASTOMA protein network controls protective quiescence in the Arabidopsis root stem cell organizer. PLoS Biol 11: e1001724

De Veylder L, Beeckman T, Beemster GT, de Almeida Engler J, Ormenese S, Maes S, Naudts M, Van Der Schueren E, Jacqmard A, Engler G, Inze D (2002) Control of proliferation, endoreduplication and differentiation by the Arabidopsis E2Fa-DPa transcription factor. EMBO J 21: 1360–8

del Pozo JC, Diaz-Trivino S, Cisneros N, Gutierrez C (2006) The balance between cell division and endoreplication depends on E2FC-DPB, transcription factors regulated by the ubiquitin-SCFSKP2A pathway in Arabidopsis. Plant Cell 18: 2224–35

Desvoyes B, Gutierrez C (2020) Roles of plant retinoblastoma protein: cell cycle and beyond. EMBO J 39: e105802

Desvoyes B, Ramirez-Parra E, Xie Q, Chua NH, Gutierrez C (2006) Cell type-specific role of the retinoblastoma/E2F pathway during Arabidopsis leaf development. Plant Physiol 140: 67–80

Dewitte W, Riou-Khamlichi C, Scofield S, Healy JM, Jacqmard A, Kilby NJ, Murray JA (2003) Altered cell cycle distribution, hyperplasia, and inhibited differentiation in Arabidopsis caused by the D-type cyclin CYCD3. Plant Cell 15: 79–92

Dong J, MacAlister CA, Bergmann DC (2009) BASL controls asymmetric cell division in Arabidopsis. Cell 137: 1320–30

Du Z, Zhou X, Ling Y, Zhang Z, Su Z (2010) agriGO: a GO analysis toolkit for the agricultural community. Nucleic Acids Res 38: W64–70

Ebel C, Mariconti L, Gruissem W (2004) Plant retinoblastoma homologues control nuclear proliferation in the female gametophyte. Nature 429: 776–80

Gaspar JM (2018) Improved peak-calling with MACS2. bioRxiv

Gutzat R, Borghi L, Futterer J, Bischof S, Laizet Y, Hennig L, Feil R, Lunn J, Gruissem W (2011) RETINOBLASTOMA-RELATED PROTEIN controls the transition to autotrophic plant development. Development 138: 2977–86

Haga N, Kobayashi K, Suzuki T, Maeo K, Kubo M, Ohtani M, Mitsuda N, Demura T, Nakamura K, Jurgens G, Ito M (2011) Mutations in MYB3R1 and MYB3R4 cause pleiotropic developmental defects and preferential down-regulation of multiple G2/M-specific genes in Arabidopsis. Plant Physiol 157: 706–17

Heyman J, Van den Daele H, De Wit K, Boudolf V, Berckmans B, Verkest A, Alvim Kamei CL, De Jaeger G, Koncz C, De Veylder L (2011) Arabidopsis ULTRAVIOLET-B-INSENSITIVE4 maintains cell division activity by temporal inhibition of the anaphase-promoting complex/cyclosome. Plant Cell 23: 4394–410

Horvath BM, Kourova H, Nagy S, Nemeth E, Magyar Z, Papdi C, Ahmad Z, Sanchez-Perez GF, Perilli S, Blilou I, Pettko-Szandtner A, Darula Z, Meszaros T, Binarova P, Bogre L, Scheres B (2017) Arabidopsis RETINOBLASTOMA RELATED directly regulates DNA damage responses through functions beyond cell cycle control. EMBO J 36: 1261–1278

Jaakola L, Pirttila AM, Halonen M, Hohtola A (2001) Isolation of high quality RNA from bilberry (Vaccinium myrtillus L.) fruit. Mol Biotechnol 19: 201–3

Kallai BM, Kourova H, Chumova J, Papdi C, Trogelova L, Kofronova O, Hozak P, Filimonenko V, Meszaros T, Magyar Z, Bogre L, Binarova P (2020) gamma-Tubulin interacts with E2F transcription factors to regulate proliferation and endocycling in Arabidopsis. J Exp Bot 71: 1265–1277

Kent LN, Leone G (2019) The broken cycle: E2F dysfunction in cancer. Nat Rev Cancer 19: 326–338

Kobayashi K, Suzuki T, Iwata E, Nakamichi N, Suzuki T, Chen P, Ohtani M, Ishida T, Hosoya H, Muller S, Leviczky T, Pettko-Szandtner A, Darula Z, Iwamoto A, Nomoto M, Tada Y, Higashiyama T, Demura T, Doonan JH, Hauser MT et al. (2015) Transcriptional repression by MYB3R proteins regulates plant organ growth. EMBO J 34: 1992–2007

Kosugi S, Ohashi Y (2002) Interaction of the Arabidopsis E2F and DP proteins confers their concomitant nuclear translocation and transactivation. Plant Physiol 128: 833–43

Lammens T, Li J, Leone G, De Veylder L (2009) Atypical E2Fs: new players in the E2F transcription factor family. Trends Cell Biol 19: 111–8

Lang L, Pettko-Szandtner A, Tuncay Elbasi H, Takatsuka H, Nomoto Y, Zaki A, Dorokhov S, De Jaeger G, Eeckhout D, Ito M, Magyar Z, Bogre L, Heese M, Schnittger A (2021) The DREAM complex represses growth in response to DNA damage in Arabidopsis. Life Sci Alliance 4

Langmead B, Salzberg SL (2012) Fast gapped-read alignment with Bowtie 2. Nat Methods 9: 357–9

Langmead B, Trapnell C, Pop M, Salzberg SL (2009) Ultrafast and memory-efficient alignment of short DNA sequences to the human genome. Genome Biol 10: R25

Legesse-Miller A, Raitman I, Haley EM, Liao A, Sun LL, Wang DJ, Krishnan N, Lemons JM, Suh EJ, Johnson EL, Lund BA, Coller HA (2012) Quiescent fibroblasts are protected from proteasome inhibition-mediated toxicity. Mol Biol Cell 23: 3566–81

Leviczky T, Molnar E, Papdi C, Oszi E, Horvath GV, Vizler C, Nagy V, Pauk J, Bogre L, Magyar Z (2019) E2FA and E2FB transcription factors coordinate cell proliferation with seed maturation. Development 146

Li X, Cai W, Liu Y, Li H, Fu L, Liu Z, Xu L, Liu H, Xu T, Xiong Y (2017) Differential TOR activation and cell proliferation in Arabidopsis root and shoot apexes. Proc Natl Acad Sci U S A 114: 2765–2770

Magyar Z, Atanassova A, De Veylder L, Rombauts S, Inze D (2000) Characterization of two distinct DP-related genes from Arabidopsis thaliana. FEBS Lett 486: 79–87

Magyar Z, Bogre L, Ito M (2016) DREAMs make plant cells to cycle or to become quiescent. Curr Opin Plant Biol 34: 100–106

Magyar Z, De Veylder L, Atanassova A, Bako L, Inze D, Bogre L (2005) The role of the Arabidopsis E2FB transcription factor in regulating auxin-dependent cell division. Plant Cell 17: 2527–41

Magyar Z, Horvath B, Khan S, Mohammed B, Henriques R, De Veylder L, Bako L, Scheres B, Bogre L (2012) Arabidopsis E2FA stimulates proliferation and endocycle separately through RBR-bound and RBR-free complexes. EMBO J 31: 1480–93

Marescal O, Cheeseman IM (2020) Cellular Mechanisms and Regulation of Quiescence. Dev Cell 55: 259–271

Mariconti L, Pellegrini B, Cantoni R, Stevens R, Bergounioux C, Cella R, Albani D (2002) The E2F family of transcription factors from Arabidopsis thaliana. Novel and conserved components of the retinoblastoma/E2F pathway in plants. J Biol Chem 277: 9911–9

Martins RP, Gandini LG, Jr., Martins IP, Martins LP (2011) Crimpable double tubes for segmental retraction. Orthodontics (Chic) 12: 400–3

Matos JL, Lau OS, Hachez C, Cruz-Ramirez A, Scheres B, Bergmann DC (2014) Irreversible fate commitment in the Arabidopsis stomatal lineage requires a FAMA and RETINOBLASTOMA-RELATED module. Elife 3

Morgan DO (2007) The Cell Cycle: Principles of Control. New Science Press,

O’Malley RC, Huang SC, Song L, Lewsey MG, Bartlett A, Nery JR, Galli M, Gallavotti A, Ecker JR (2016) Cistrome and Epicistrome Features Shape the Regulatory DNA Landscape. Cell 165: 1280–1292

Oszi E, Papdi C, Mohammed B, Petko-Szandtner A, Leviczky T, Molnar E, Galvan-Ampudia C, Khan S, Juez EL, Horvath B, Bogre L, Magyar Z (2020) E2FB Interacts with RETINOBLASTOMA RELATED and Regulates Cell Proliferation during Leaf Development. Plant Physiol 182: 518–533

Ramirez F, Ryan DP, Gruning B, Bhardwaj V, Kilpert F, Richter AS, Heyne S, Dundar F, Manke T (2016) deepTools2: a next generation web server for deep-sequencing data analysis. Nucleic Acids Res 44: W160–5

Robinson MD, McCarthy DJ, Smyth GK (2010) edgeR: a Bioconductor package for differential expression analysis of digital gene expression data. Bioinformatics 26: 139–40

Scheres B (2007) Stem-cell niches: nursery rhymes across kingdoms. Nat Rev Mol Cell Biol 8: 345–54

Schneider CA, Rasband WS, Eliceiri KW (2012) NIH Image to ImageJ: 25 years of image analysis. Nat Methods 9: 671–5

Shimada T, Fuji K, Tamura K, Kondo M, Nishimura M, Hara-Nishimura I (2003) Vacuolar sorting receptor for seed storage proteins in Arabidopsis thaliana. Proc Natl Acad Sci U S A 100: 16095–100

Sozzani R, Maggio C, Varotto S, Canova S, Bergounioux C, Albani D, Cella R (2006) Interplay between Arabidopsis activating factors E2Fb and E2Fa in cell cycle progression and development. Plant Physiol 140: 1355–66

Umbrasaite J, Schweighofer A, Kazanaviciute V, Magyar Z, Ayatollahi Z, Unterwurzacher V, Choopayak C, Boniecka J, Murray JA, Bogre L, Meskiene I (2010) MAPK phosphatase AP2C3 induces ectopic proliferation of epidermal cells leading to stomata development in Arabidopsis. PLoS One 5: e15357

Vandepoele K, Raes J, De Veylder L, Rouze P, Rombauts S, Inze D (2002) Genome-wide analysis of core cell cycle genes in Arabidopsis. Plant Cell 14: 903–16

Vanstraelen M, Baloban M, Da Ines O, Cultrone A, Lammens T, Boudolf V, Brown SC, De Veylder L, Mergaert P, Kondorosi E (2009) APC/C-CCS52A complexes control meristem maintenance in the Arabidopsis root. Proc Natl Acad Sci U S A 106: 11806–11

Verkest A, Abeel T, Heyndrickx KS, Van Leene J, Lanz C, Van De Slijke E, De Winne N, Eeckhout D, Persiau G, Van Breusegem F, Inze D, Vandepoele K, De Jaeger G (2014) A generic tool for transcription factor target gene discovery in Arabidopsis cell suspension cultures based on tandem chromatin affinity purification. Plant Physiol 164: 1122–33

Wang S, Gu Y, Zebell SG, Anderson LK, Wang W, Mohan R, Dong X (2014) A noncanonical role for the CKI-RB-E2F cell-cycle signaling pathway in plant effector-triggered immunity. Cell Host Microbe 16: 787–94

Wildwater M, Campilho A, Perez-Perez JM, Heidstra R, Blilou I, Korthout H, Chatterjee J, Mariconti L, Gruissem W, Scheres B (2005) The RETINOBLASTOMA-RELATED gene regulates stem cell maintenance in Arabidopsis roots. Cell 123: 1337–49

Wyrzykowska J, Schorderet M, Pien S, Gruissem W, Fleming AJ (2006) Induction of differentiation in the shoot apical meristem by transient overexpression of a retinoblastoma-related protein. Plant Physiol 141: 1338–48

Xiang G, Keller CA, Giardine B, An L, Li Q, Zhang Y, Hardison RC (2020) S3norm: simultaneous normalization of sequencing depth and signal-to-noise ratio in epigenomic data. Nucleic Acids Res 48: e43

Yao G (2014) Modelling mammalian cellular quiescence. Interface Focus 4: 20130074

Yao G, Tan C, West M, Nevins JR, You L (2011) Origin of bistability underlying mammalian cell cycle entry. Mol Syst Biol 7: 485

Yao X, Yang H, Zhu Y, Xue J, Wang T, Song T, Yang Z, Wang S (2018) The Canonical E2Fs Are Required for Germline Development in Arabidopsis. Front Plant Sci 9: 638

Zhao X, Bramsiepe J, Van Durme M, Komaki S, Prusicki MA, Maruyama D, Forner J, Medzihradszky A, Wijnker E, Harashima H, Lu Y, Schmidt A, Guthorl D, Logrono RS, Guan Y, Pochon G, Grossniklaus U, Laux T, Higashiyama T, Lohmann JU et al. (2017) RETINOBLASTOMA RELATED1 mediates germline entry in Arabidopsis. Science 356

