## supplemental figures and table for "The canonical E2Fs together with RETINOBLASTOMA-RELATED are required to establish quiescence during plant development"

### Extended Data Fig. 1

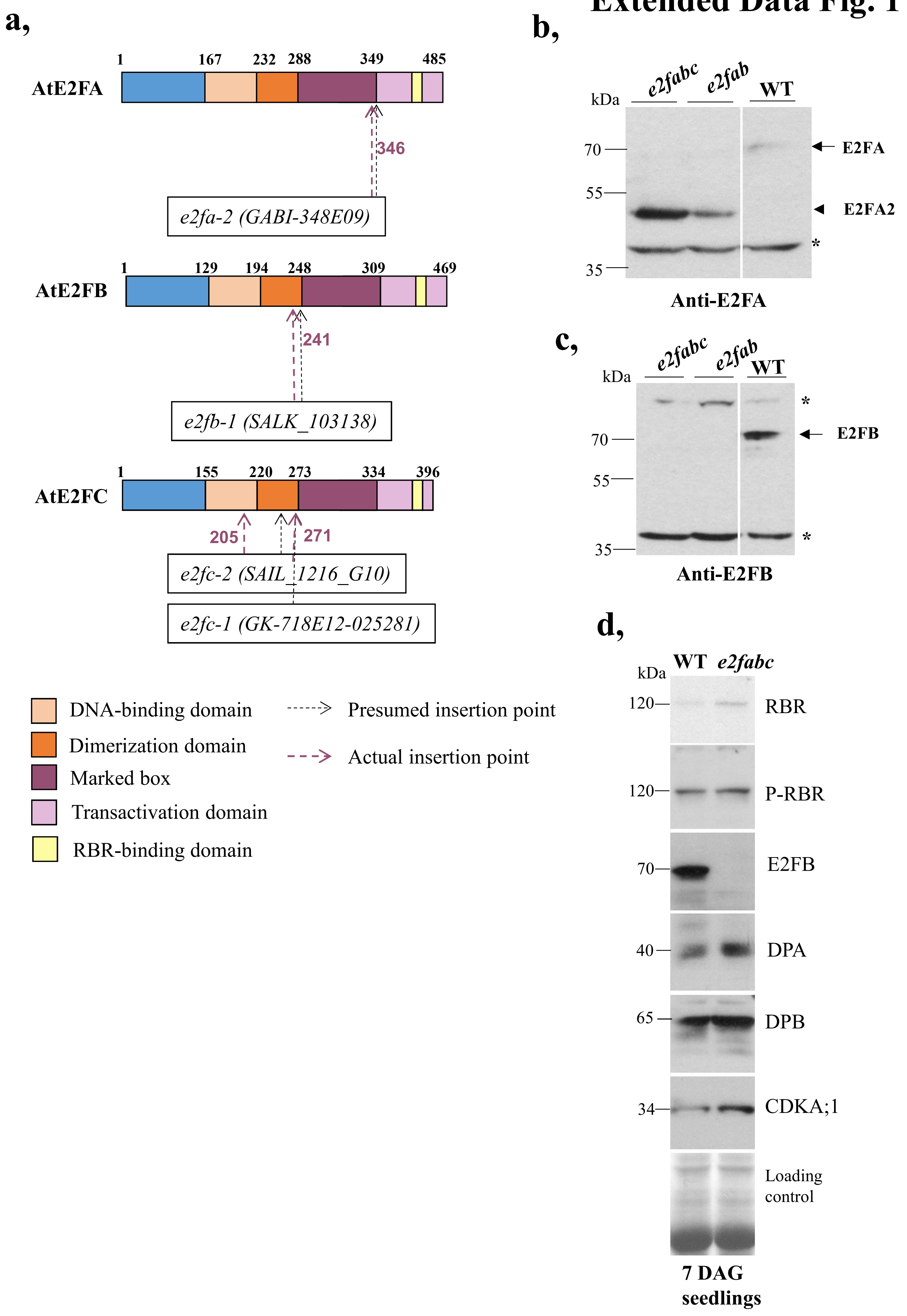

**Extended Data Figure 1: Characterization of E2F mutant lines used in this work.**

a: Position of the T-DNA insertions in mutant lines used in this study. In each mutant the exact position of the insertion was verified by sequencing the flanking regions and the presumed and the actual insertion points are shown by black and purple arrows, respectively. For all genes, the regions encoding each protein domain are highlighted with different colours.

b, c: Accumulation of E2FB is undetectable in *e2fab*, and *e2fabc* mutants using a C-terminal specific anti-E2FB antibody in western blot whereas the N-terminal specific anti-E2FA antibody recognized a truncated version of the E2FA protein in the same *e2f* mutants.

d: The accumulation level of RBR, phosphorylated RBR (P-RBR$^{911Ser}$), CDKA;1, DPA and DPB proteins was comparable in the *e2fabc* with the WT or even slightly enhanced (description of these antibodies are found in the Methods section). Seedlings at 7DAG were used in Western blot assay.

### Extended Data Fig. 2

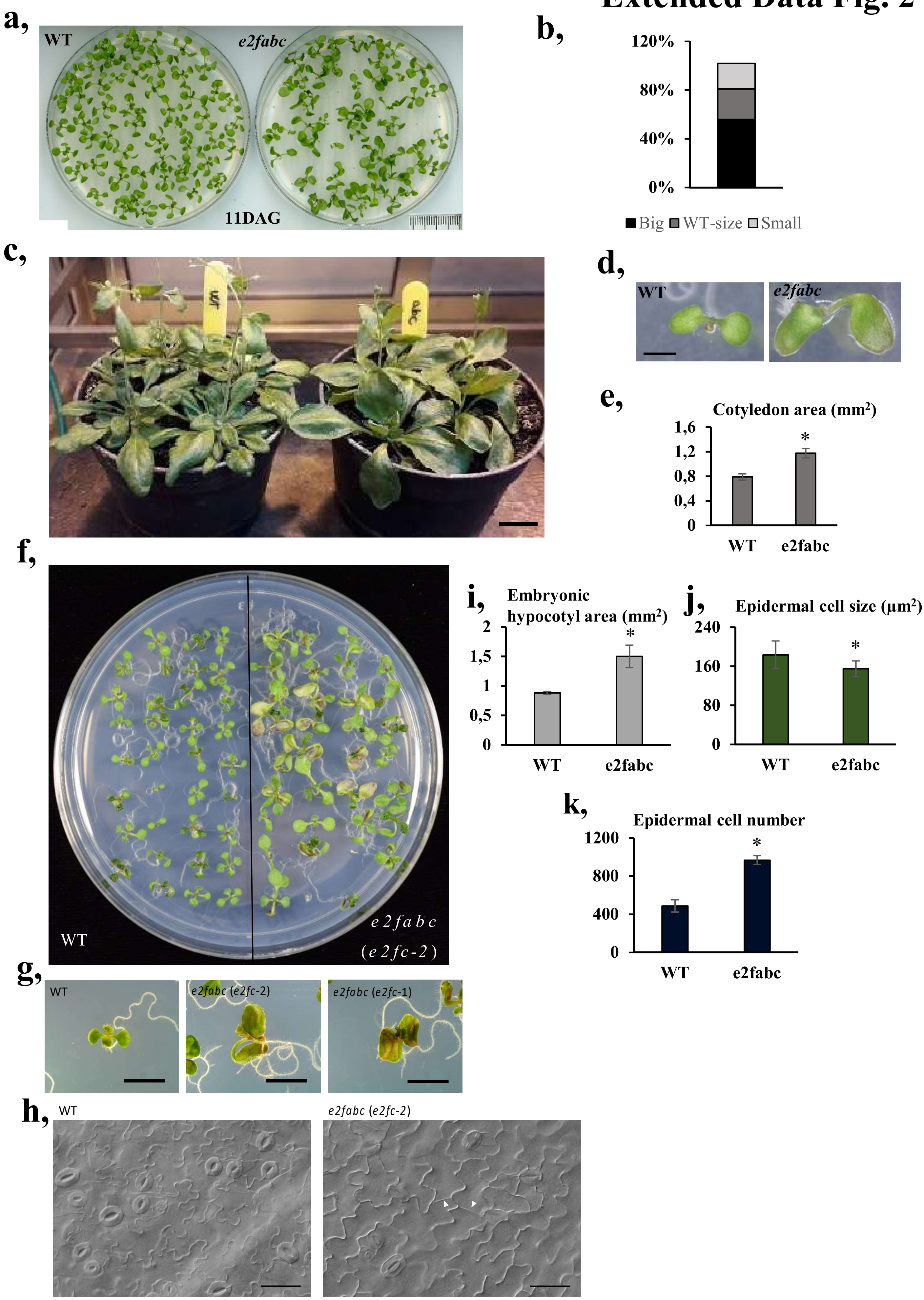

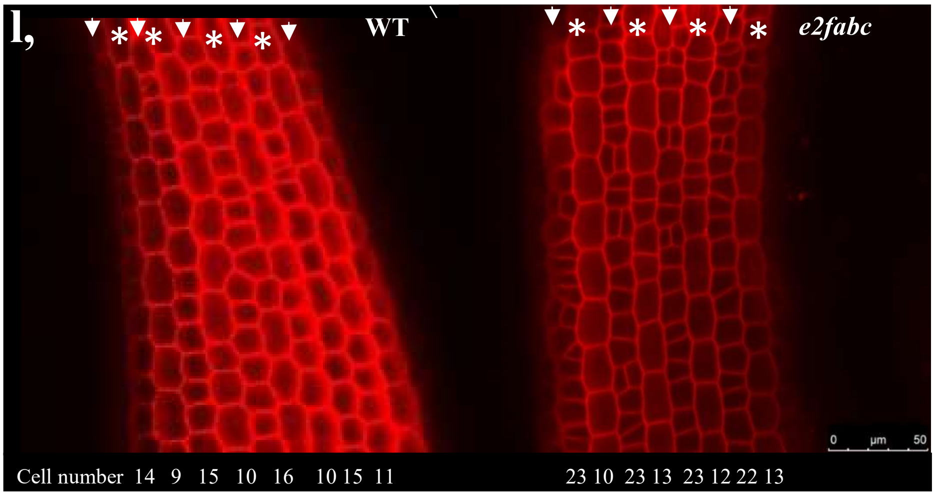


**External Data Figure 2: *e2fabc* mutants are enlarged compared to the WT.**

a: Representative phenotypes of plantlets at 11 DAG grown on agar medium.

b: Distribution of *e2fabc* plantlets between big, WT-size and small categories.

c: Picture of WT and *e2fabc* plants with fully expanded rosettes grown for 23 days on soil.

d,e: Cotyledons of *e2fabc* mutants are enlarged. d: Pictures of plantlets at 7 DAG, e: Graph showing cotyledon area in plantlets at 7 DAG. Data are average +/- standard deviation (n=3 biological replicates, N=10 samples in each). *$P \leq 0.05$, denotes statistically relevant differences (two-tailed, paired *t*-test between the WT and the mutant).

f-h: Young seedlings of *e2fabc-2* triple mutant are enlarged. f: Pictures of plantlets at 9 DAG, g: Higher magnification of seedlings at 9 DAG, scale bar = 5 mm, h: DIC image of differentiated pavement cells in the leaf epidermis of the triple *e2fabc-2* line show newly synthesised cell walls indicating cell proliferation. Scale bars = 20 µm.

i-l: Hypocotyls of *e2fabc* embryos are enlarged and consist of more numerous and smaller cells compared to the WT. f: Embryonic hypocotyl area, g: Epidermal cell size, h: Epidermal cell number, i: Representative confocal microscopy image showing the hypocotyl epidermis after PI staining. Asterisks show longitudinal cell files where extra cell divisions were observed in the *e2fabc* mutant but not in the WT. For all graphs, data are average +/- standard deviation, (n=3 biological repeats, N=10 samples in each), *$P \leq 0.05$, **$P \leq 0.01$ denotes statistically relevant differences (two-tailed, paired *t*-test between the WT and the mutant).

### Extended Data Fig. 3

a,

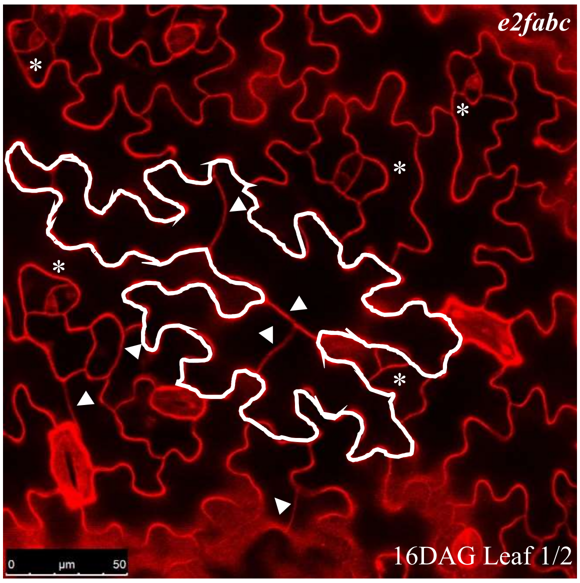


b,

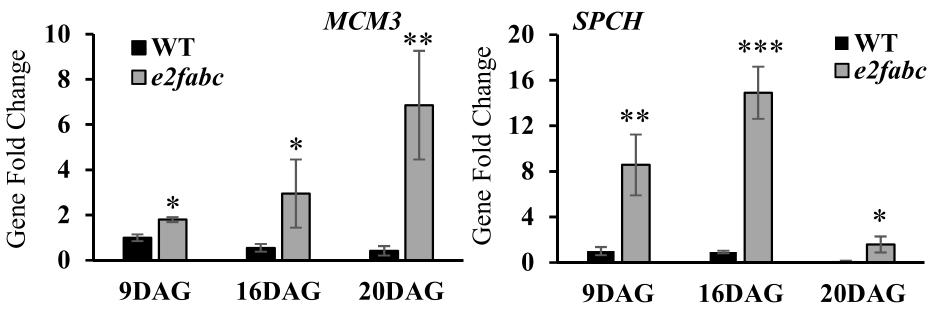


**External Data Figure 3: Cell proliferation is enhanced in *e2fabc* mutants.**

a: *e2fabc* mutant leaves display extra cell division events in puzzle-shaped differentiated epidermal cells.

b, c: The cell cycle gene *MCM3* and the stomata development gene *SPCH* are over-expressed in the first leaf pairs of *e2fabc* mutants compared to the WT at 9, 16, 20 DAG. Expression of the two genes was quantified by qRT-PCR, and values represent fold change levels normalised to the relevant transcript levels of the WT at 9DAG, which was set arbitrarily at 1. n=3 biological repeats. Error bars indicate the +/- SD. *$P \leq 0.05$, **$P \leq 0.01$, ***$P \leq$ 0.001; indicate statistical significance two-tailed, paired *t*-test between the wild type and the e2fabc at a given time point. Abbreviations and primer sequences are listed in External Data Table 1.

#### Extended Data Fig. 4

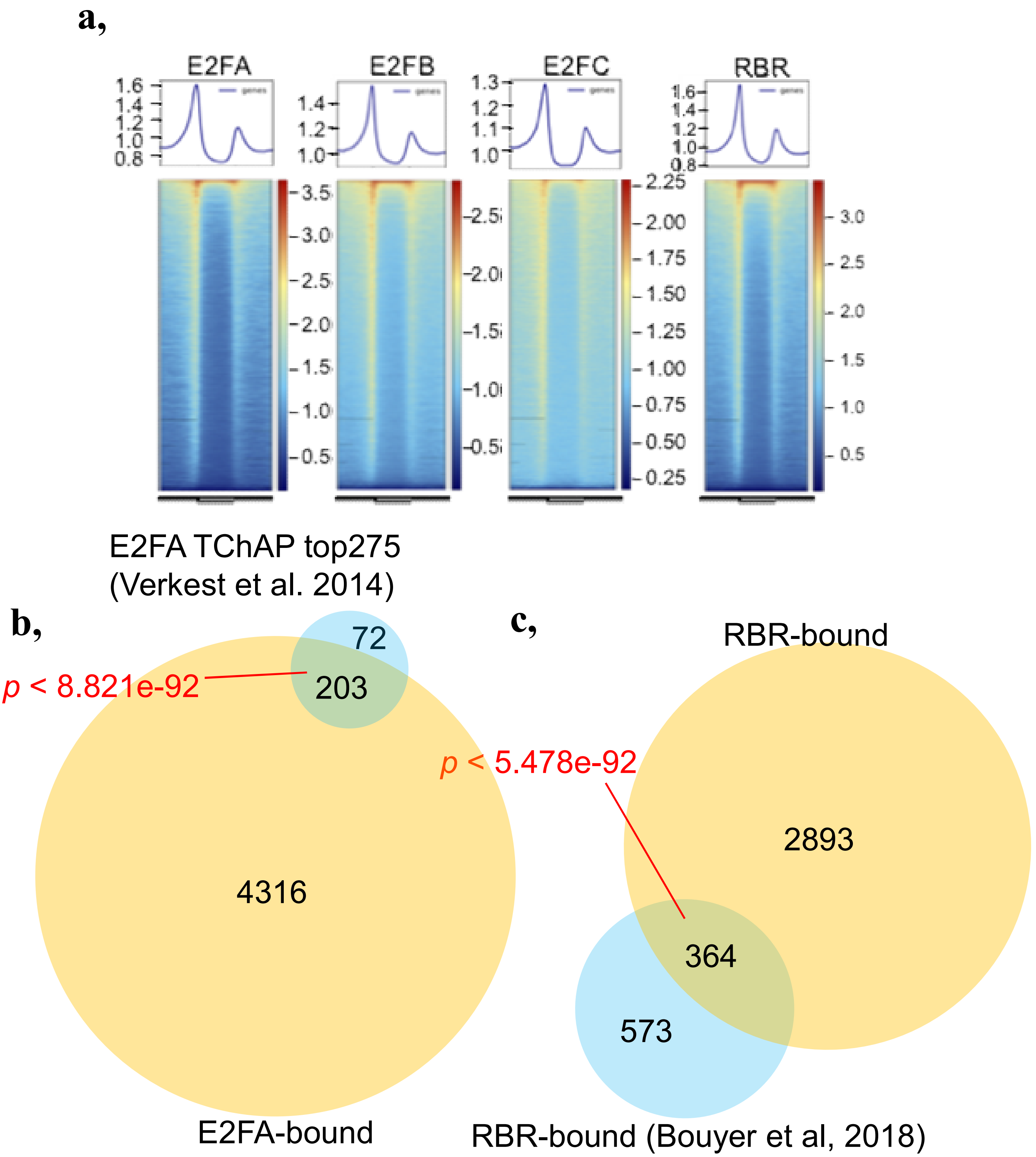


**External Data Figure 4: Quality control of the ChIP-seq analysis**

a: Metaplot and heatmap showing the binding profiles of E2FA, E2FB, E2FC and RBR. b: comparison of E2FA targets identified in this work, and the top 275 genes defined in previous TChAP experiments (Verkest et al., 2014). c: comparison of RBR targets identified in this work, and those targets reported by (Bouyer, Heese et al., 2018). For both comparisons, overlaps obtained are significantly greater than what would be expected by chance, *P*-values are indicated in red (Fisher exact test).

### Extended Data Fig. 5

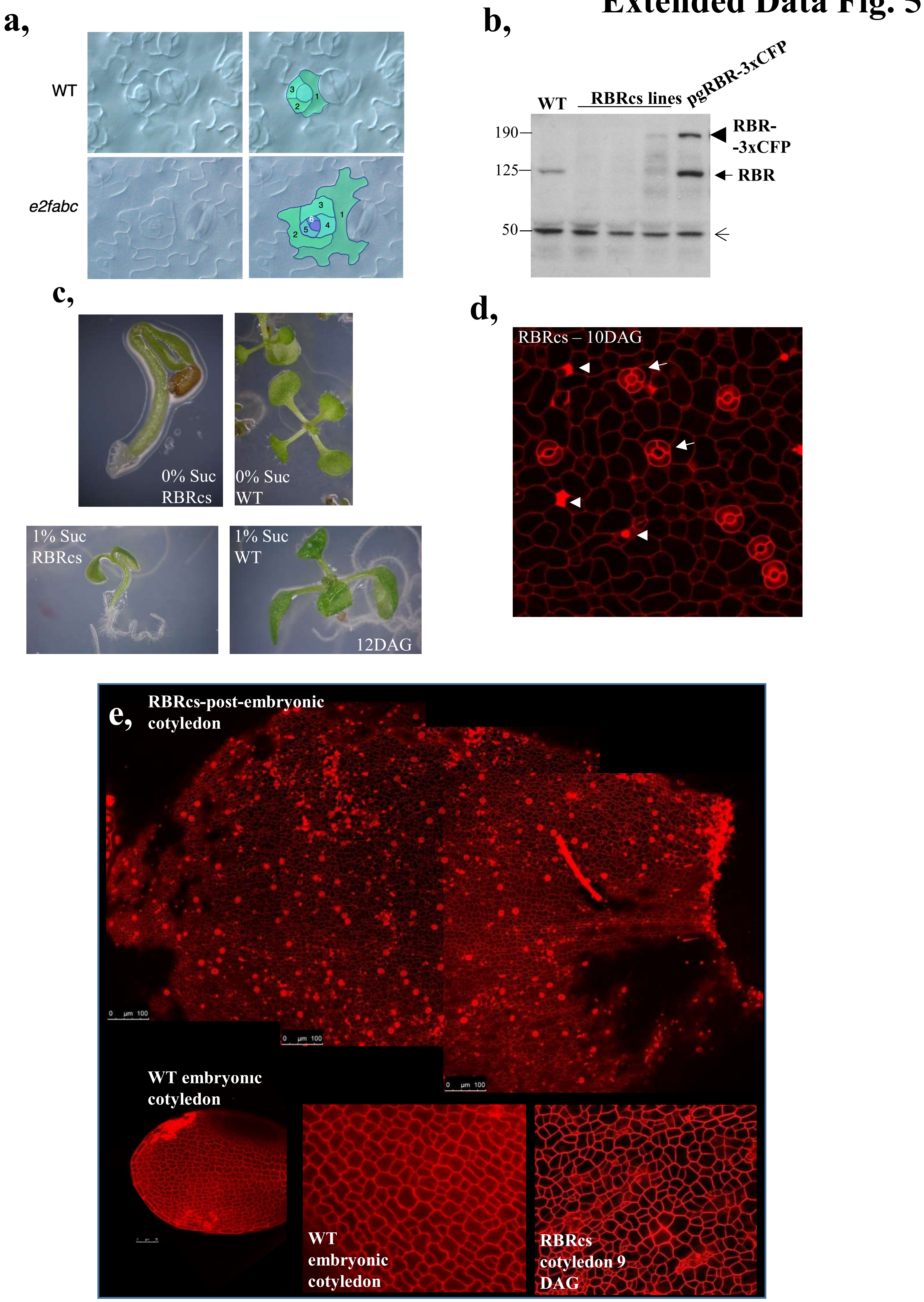

**External Data Figure 5: Comparison of cellular phenotypes observed in the epidermis of *e2fabc* and RBRcs lines.**

a: DIC image of a stomata meristemoid (top) and cartoon highlighting reiterated cell division events.

b: Immunoblot analysis of WT, RBR-CFP and *RBRcs* seedlings 6 DAG. Arrowhead shows RBR-3x-CFP, arrow indicates RBR proteins, open arrowhead marks a protein that cross-reacted with RBR antibody and is present in each lane at comparable levels.

c: All mutant seedlings with reduced RBR levels had identical phenotypic abnormalities as reported earlier (Gutzat et al., 2011). On sucrose free medium the root of the *RBRcs* seedlings did not grow, and cotyledons were closed (12DAG), while in the presence of sucrose roots grow and cotyledons were open but they never produced expanding first leaf pairs, and their growth was strongly arrested in comparison to the WT at the same age.

d-e: PI stained cotyledons of RBRcs seedlings (at 9 DAG) and WT embryos. The obtained images are highly similar, showing that epidermal cells of RBRcs remain embryonic-like. Arrows show stomata consisting of three cells, while arrowheads indicate dead cells in d.

### Extended Data Fig. 6

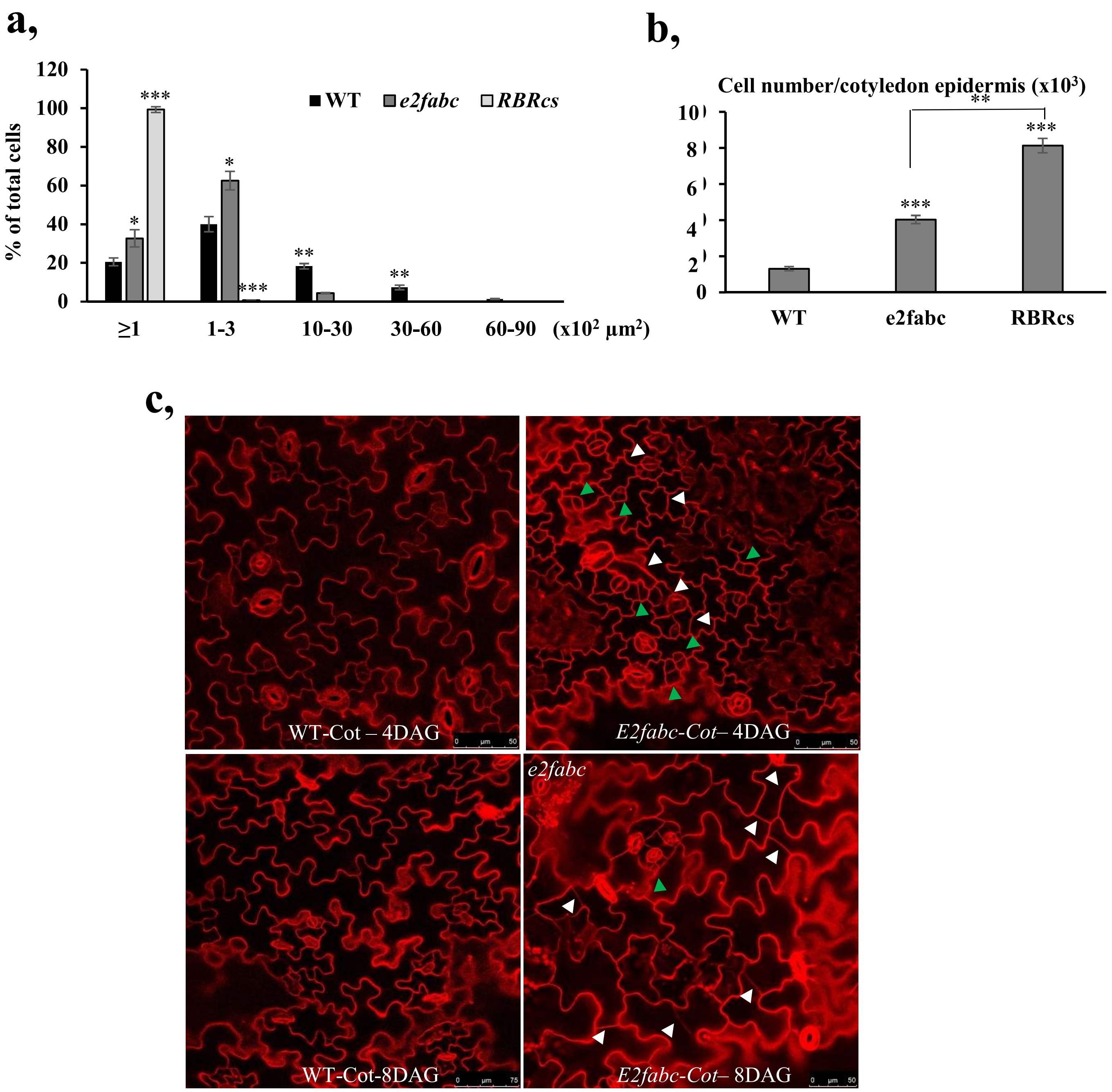


**External Data Figure 6: Cell proliferation is enhanced in *e2fabc* mutants and RBRcs lines**

a: Cell size distribution in the cotyledon epidermis of *e2fabc* and RBRcs lines is shifted towards the smaller cell sizes. n=3 biological repeats, N=10 samples in each. ≥400 cells were measured using ImageJ. **P*<0.05, ***P*≤0.01, ****P*≤0.001, significant difference in the transgenic lines compared to the WT using Student t-test.

b: Cell number is increased in the cotyledon epidermis of RBRcs lines and to a lesser extent in *e2fabc* lines. ***P*≤0.01, ****P*≤0.001; indicate statistical significance (two-tailed, paired *t*-test between the WT and the mutants, and between the two mutants).

c: Representative images of PI stained cotyledon epidermis of WT and *e2fabc* mutants. White arrowheads point at clustered meristemoids, and green arrowheads point at extra cell division events occurring in differentiated cells.

### Extended Data Fig. 7

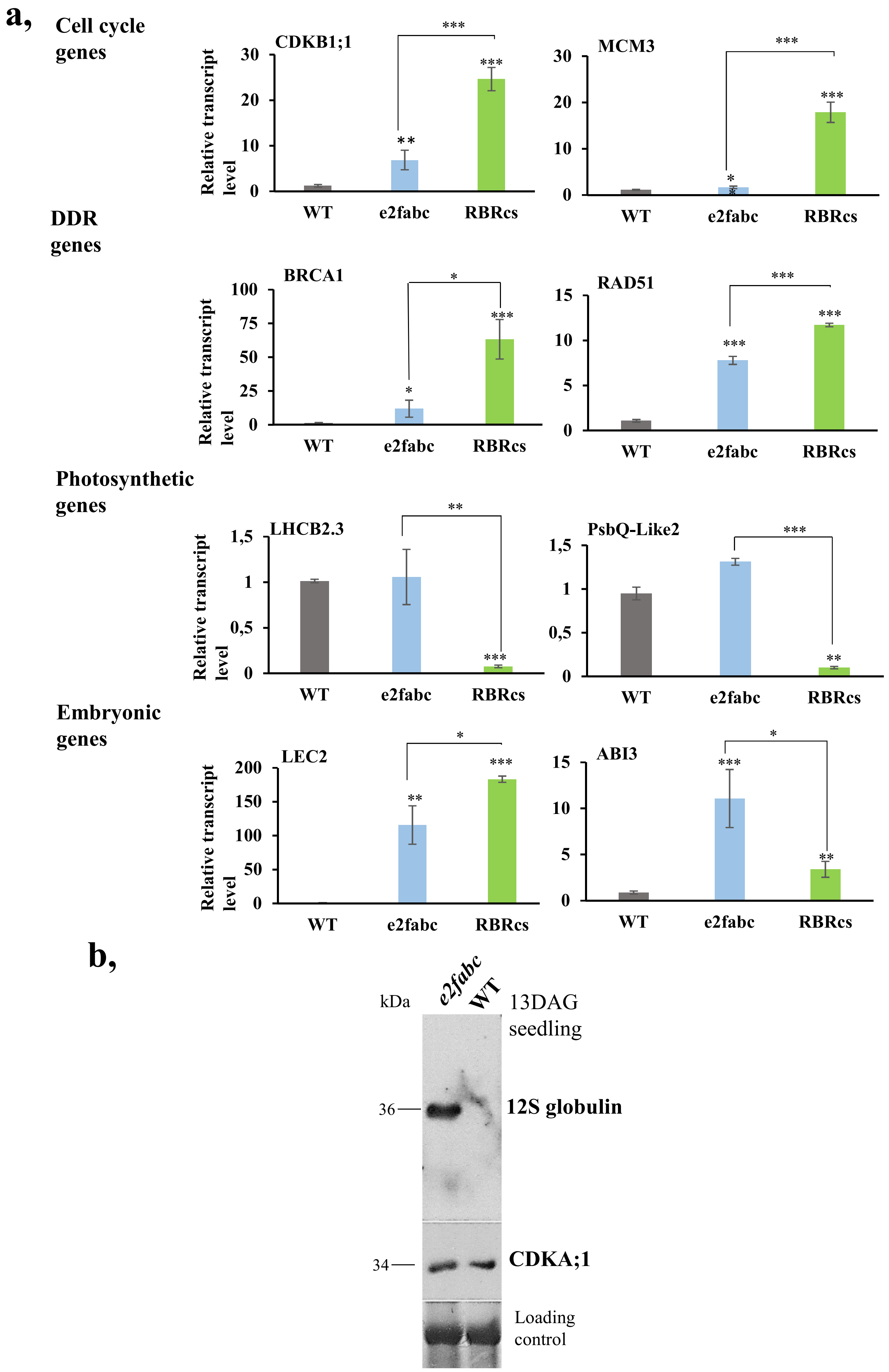

**External Data Figure 7: RT-qPCR quantification of marker genes in *e2fabc* and *RBRcs* lines**

a, Cell cycle and DNA repair genes are highly induced in *RBRcs* lines, and to a lesser extent in *e2fabc* mutants. By contrast, differentiation-related genes are specifically repressed in RBRcs, but not in *e2fabc* mutants. Values represent fold change levels normalised to the relevant transcript levels in the WT, which was set arbitrarily at 1. n=3 biological repeats. Error bars indicate the SD. *$P$<0.05, **$P$≤0.01, ***$P$≤0.001 indicates statistical significance determined using two-tailed, paired *t*-test between the WT and the mutants and between the mutants. Abbreviations and primer sequences are listed in External Data Table 1. b, Seed storage 12S globulin accumulated in the *e2fabc* mutant seedlings 13 DAG but not in the WT at the same age. CDKA;1 proteins were at comparable levels in both of these lines. Coomassie stained proteins were used as loading control.

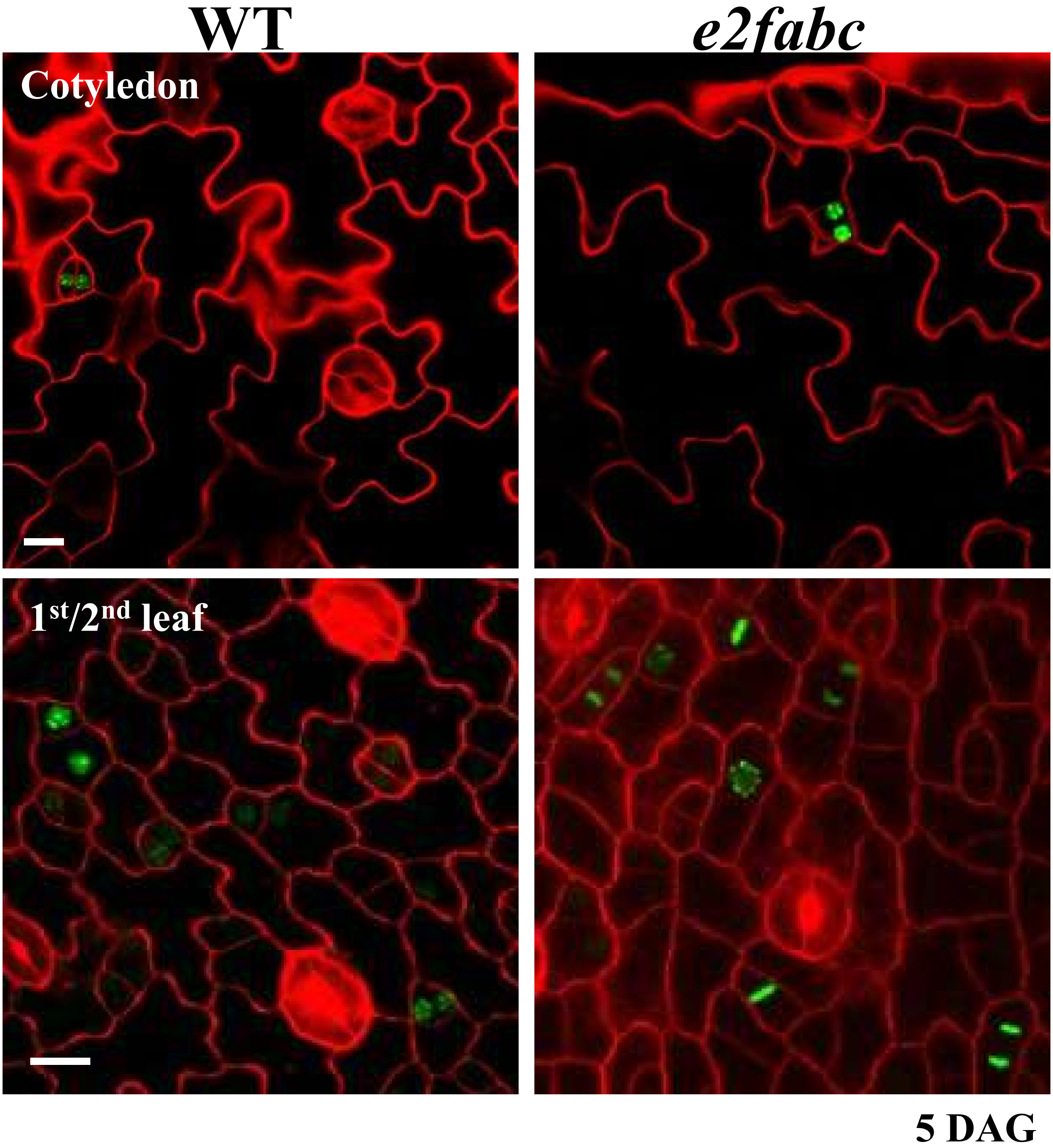


**External Data Figure 8: Expression of the cell division marker CYCB1;2 is enhanced in *e2fabc* mutants.**

Cotyledons and first leaves of WT and e2fabc mutants expressing a YFP-tagged version of CYCB1;2 were stained with PI. Expression of the cell division marker was low in cotyledons of both genotypes, but markedly increased in the first leaves of *e2fabc* mutants compared to the WT. Scale bars = 10 µm.

### Extended data Fig. 9

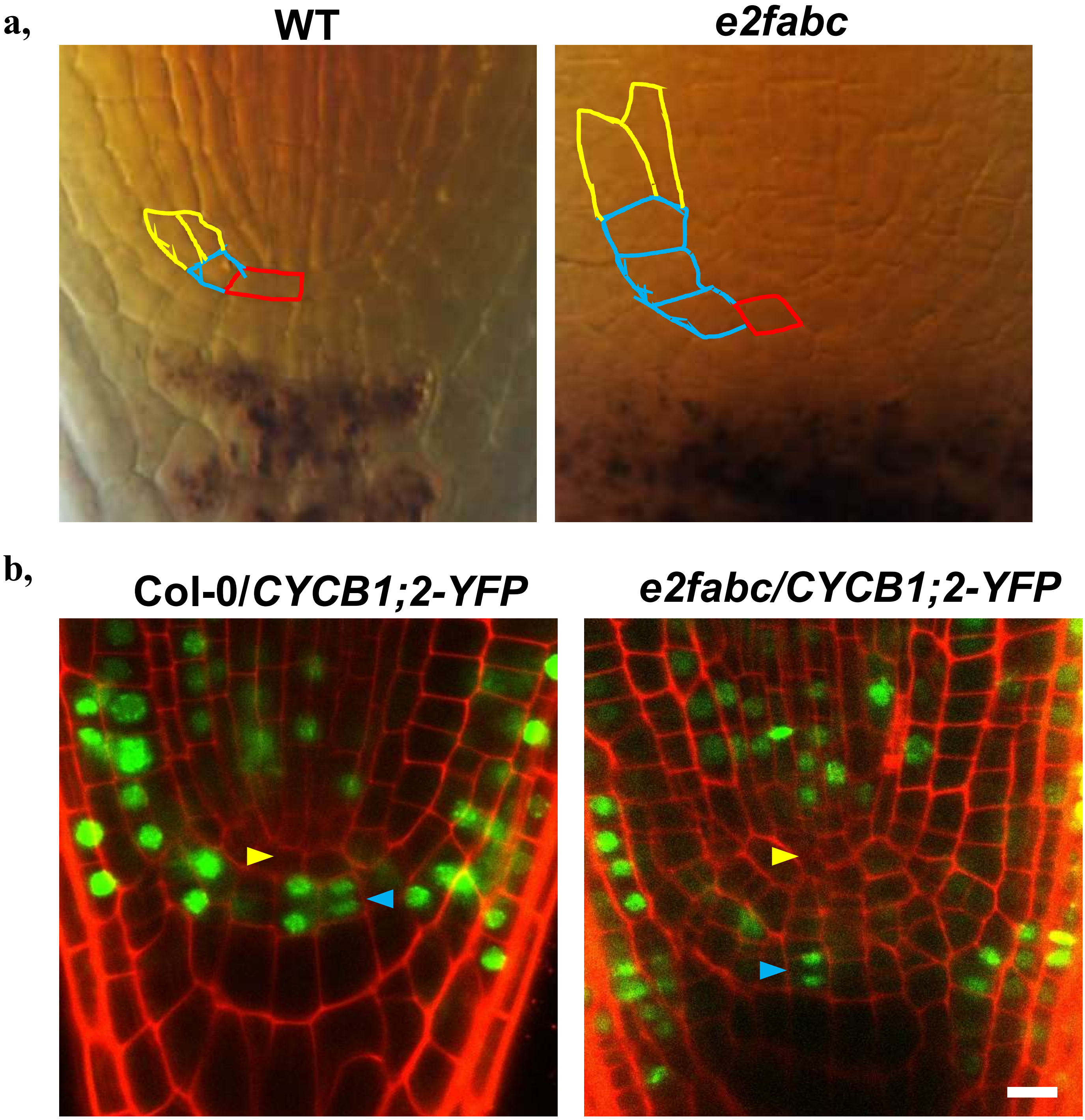


**External Data Figure 9: Enhanced cell proliferation of stem cells in *e2fabc* root meristems**

a: DIC images showing that *e2fabc* mutants display an increased number of cortex/endodermis initials (drawn in blue) compared to the WT. The first cortex and endodermis cells generated by asymmetric division are drawn in yellow, and the QC is drawn in red.

b: expression of the CYCB1;2-YFP marker in the root meristem of WT and *e2fabc* mutants. Blue arrowheads indicate cell division events in the columella initials. One such event is observed in supernumerary columella initials of the *e2fabc* mutant. Scale bar: 10 µm.

### Extended Data Fig. 10

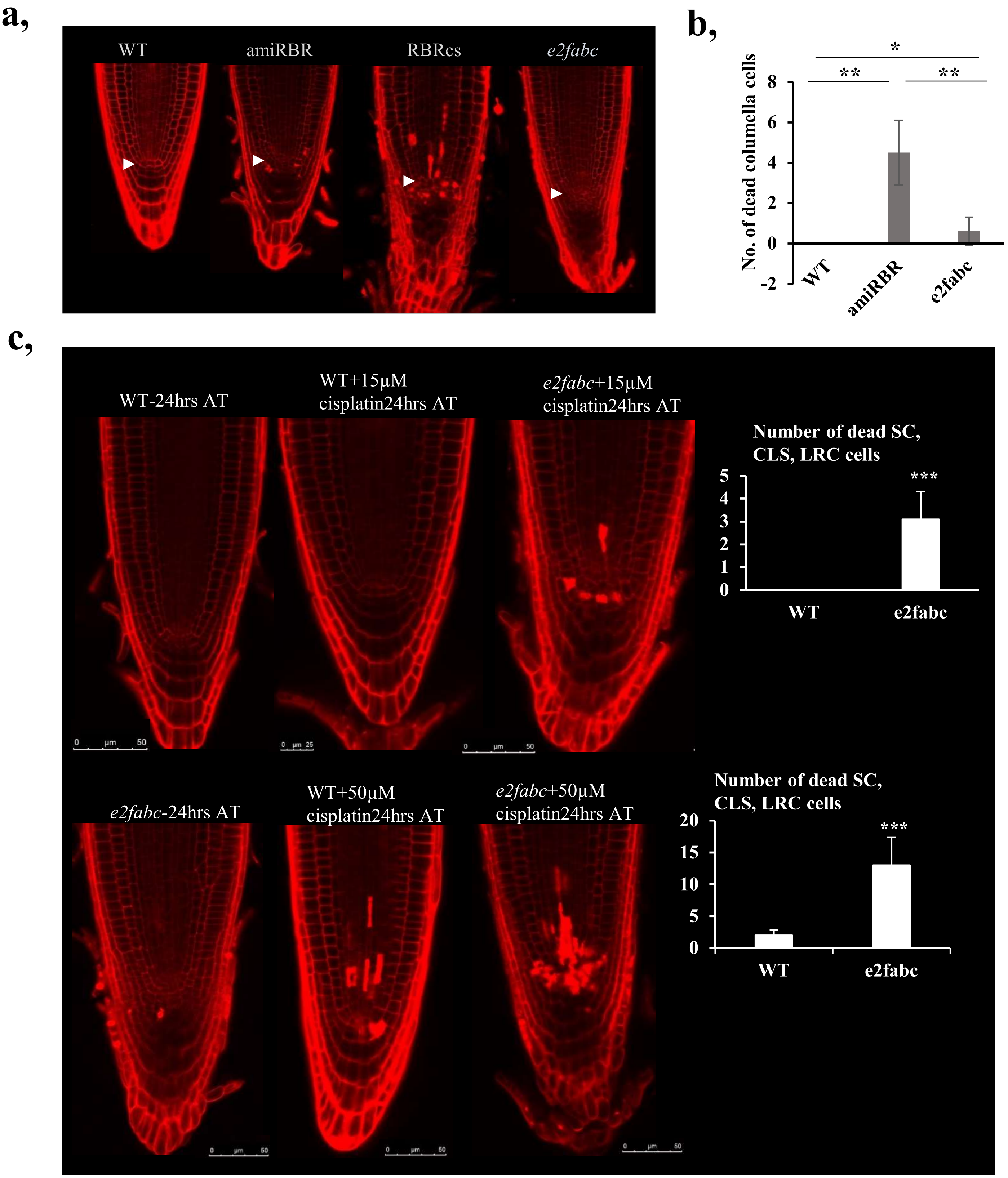


**External Data Figure 10: Loss of quiescence in RBR loss of function and *e2fabc* lines affects meristem maintenance**

a: Spontaneous cell death is observed in meristems of RBR loss of function lines but not in *e2fabc* mutants in control conditions. Root tips of seedlings at 4 DAG were stained with PI and imaged under a confocal microscope.

b: Quantification of cell death in the columella of RBRcs and *e2fabc* mutants. Data are average +/- SD, n=3 biological replicates, N=8 in each. *$P \leq 0.05$, **$P \leq 0.01$; indicate statistical significance using Student's *t*-test comparing amiRBR and *e2fabc* to WT and amiRBR to the *e2fabc*.

c: *e2fabc* mutants are hypersensitive to cisplatin. Plantlets of WT and *e2fabc* mutants at 5 DAG were treated with the indicated dose of cisplatin for 24h, and stained with PI prior to confocal imaging. Cell death was more strongly induced by both doses of cisplatin in the *e2fabc* mutant than in the WT.

### External Data Fig. 11

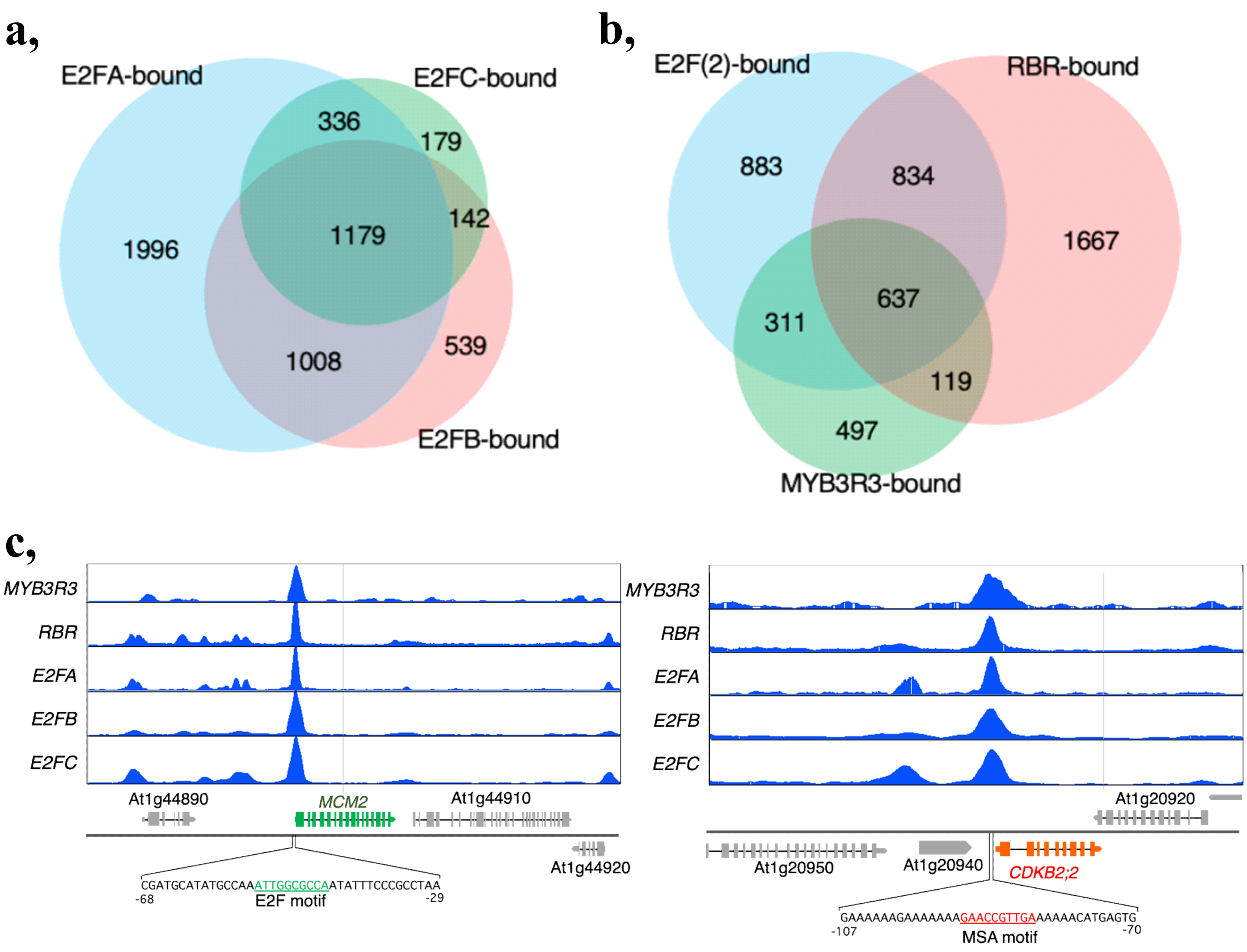


**Extended Data Fig 11: Overlap between E2F and MYB3R targets**

a: Venn diagram showing the overlap between the best targets for each E2F (FC>3 *P*<0.01).

b: Venn diagram showing the overlap between genes bound by at least two E2F factors E2F(2) and the best RBR and MYB3R3 targets (FC>3 *P*<0.01).

c: Screenshots illustrating that E2FA,B,C, RBR and MYB3R3 are found at the same position on their common targets. Here we chose MCM2, on which all factors are found in the region harbouring the canonical E2F binding site, and CDKB1;2, on which all 4 factors are found in the region harbouring the canonical MSA binding site.

| GI | NAME | FWD | REV | PROD. SIZE |
|---|---|---|---|---|
| AT1G80080 | TMM | GGACGTGAAGCATCTAAGCGA | TCAGCTTTCTCCTCATCCTCC | 101 |
| AT2G37560 | ORC2 | ACCATGTCAACGCTCCATTA | TCGACATTGTATGGTGCAAA | 100 |
| AT1G28300 | LEC2 | GCAAGAATCTCTACTTCGCC | CTTCCTCTTCGTCTCTTGGT | 129 |
| AT5G11510 | MYB3R4 | AATCGCTTGAGAAAGTAGACC | AGTAGACAGGACTGGCTTACCG | 140 |
| AT3G09370 | MYB3R3 | AGTATCACCTACTCATAGGTAC | AGCTCTTGCCTTTAAACGTGTC | 125 |
| AT3G01440 | PQL2/PnsL3 | ACAAGAACAGAGGCTGACACC | CGGTTTTACGACTCTCGGGT | 122 |
| AT3G27690 | LHCB | CAAGTCTACTCCTCAGAGCA | CGTAGTCTCCAGGGTATTCTC | 110 |
| AT4G21070 | BRCA1 | TCATGGGAGATTTCGAGCTT | ATTTAGCCAAGGCTTCAGCA | 195 |
| AT5G20850 | RAD51 var 88 &92 | TCCCTGTCGAACAGCTTCAG | CCTGTCTCTGAGCATGGAGC | 225 |
| AT3G54180 | CDKB1;1 | TCTGTTGGTTGTATCTTTGCTGA | CATTGCTGCTCAGTTGGTGT | 119 |
| AT3G24650 | ABI3 | GGCAGGGATGGAAACCAGAAAAGA | GGCAAAACGATCCTTCCGAGGTTA | 94 |
| AT5G53210 | SPCH | GCTGCTCTTGAAGATTTGGCT | CACTCAATTCCAATCTTGATGGTG | 101 |
| AT1G08560 | KNOLLE | GCTCGGATCGAACAGTACCA | CGCCTTCCTCAAACCAGACA | 111 |
| AT5G46280 | MCM3 | TGGGCAGCACATGAGGAC | CACTTTGTTATCATCTTGCAGTTT | 148 |
| AT5G42990 | UBC18 | ACAGCAATGGACATATTTGTTTAGA | TGATGCAGACTGAACTCACTGTC | 78 |
| AT3G18780 | ACTIN | GACCTTTAACTCTCCCGCTATG | CAGAATCCAGCACAATACCG | 92 |

**Extended Data Table 1: List of primers and their sequences used for qRT-PCR analysis**
